# A dissemination-prone morphotype enhances extrapulmonary organ entry by the fungus *Cryptococcus neoformans*

**DOI:** 10.1101/2020.08.27.270843

**Authors:** Steven T. Denham, Brianna Brammer, Krystal Y. Chung, Morgan A. Wambaugh, Joseph M. Bednarek, Li Guo, Jessica C.S. Brown

## Abstract

Environmental pathogens, which move from ecological niches to mammalian hosts, must adapt to dramatically different environments. Microbes that disseminate farther, including the fungal meningitis pathogen *Cryptococcus neoformans*, require additional adaptation to diverse tissues. When *C. neoformans* enters the lungs, infecting cells (<10 μm diameter) enlarge (>30 μm diameter), then form a heterogeneous population. The brain contains uniformly small cells (∼7 μm). We demonstrate that formation of a small *C. neoformans* morphotype – called “seed” cells due to their disseminating ability – is critical for extrapulmonary organ entry. Seed cell formation is triggered by environmental factors, including *C. neoformans*’ environmental niche, pigeon guano. The underlying trigger, phosphate, can be released by tissue damage, potentially establishing a feed-forward loop of seed cell formation and dissemination. We demonstrate that *C. neoformans*’ size variation is not just a continuum but inducible subpopulations that change host interactions to facilitate microbe survival and spread.

## INTRODUCTION

Environmental pathogens must be able to adapt to a wide variety of conditions as they transition from their ecological niches to host infection. This is particularly challenging for pathogens that cause disseminated disease, such as the primary cause of fungal meningitis, *Cryptococcus neoformans* (Rajasingham et al., 2017). As disseminating pathogens such as *C. neoformans* escape from the lungs, they must evade the immune system (Bojarczuk et al., 2016; Bulmer and Sans, 1967; Chun et al., 2011; Luberto et al., 2003; Okagaki et al., 2010; Stano et al., 2009; Walsh et al., 2019; Zaragoza et al., 2010), survive in the bloodstream and/or host cells (Botts and Hull, 2010; Charlier et al., 2009; Diamond and Bennett, 1973; Feldmesser et al., 2000; Gaylord et al., 2020; Walsh et al., 2019), and finally enter and grow in the brain (Chang et al., 2004; Chen et al., 2003; Chen et al., 2021; Huang et al., 2011; Liu et al., 2013; Liu et al., 2014; Maruvada et al., 2012; Olszewski et al., 2004; Santiago-Tirado et al., 2017; Vu et al., 2013).

Many microbes employ phenotypic switching and/or phenotypic heterogeneity to survive in complex and fluctuating host environments (Altamirano et al., 2020; Weigel and Dersch, 2018). These strategies include the antigen switching employed by bacterial species (spp.) including *Neisseria* spp. (Haas and Meyer, 1986; Swanson et al., 1986), *Borrelia burdorferi* (Chaconas et al., 2020), and parasites such as *Plasmodium* spp. (Roberts et al., 1992; Smith et al., 1995), which allow the infecting organism to avoid adaptive host immune responses. Among fungi, *Candida albicans* adjust exposure levels of β-glucan (Ballou et al., 2016), which is readily detected by the host (Brown et al., 2003; Gow et al., 2007).

Another strategy is phenotypic heterogeneity (Altamirano et al., 2020), a mechanism of bet-hedging against environmental stressors, including host responses that can occur within single foci of infection (Davis, 2018). For example, microbial cells at the perimeter of infection foci may be more open to attack from immune cells, while the core may be depleted of nutrients and oxygen (Davis et al., 2015; Kowalski et al., 2019). Variation within populations provide phenotypic robustness and can also influence which cells in a localized population are more likely to spread to new sites within a host, a phenomenon that significantly impacts the likelihood and trajectory of cancer metastasis (San Juan et al., 2019). Fungi have long been known to undergo dramatic phenotypic changes that influence disease, such as the yeast / hyphal switches of dimorphic fungi (Garfoot et al., 2016; Guimarães et al., 2011; Kanetsuna and Carbonell, 1971; Mukaremera et al., 2017; Viriyakosol et al., 2013; Whiston et al., 2012), spherule development in *Coccidioides* spp. (Viriyakosol et al., 2013; Whiston et al., 2012), and sporulation of conidia in *Aspergillus* spp. (Aimanianda et al., 2009; Hohl et al., 2005). The differences between yeast and filamentous forms are dramatic and superficially obvious due to their extreme morphological differences. However, we are only beginning to understand that more subtle variations within morphological states can have profound influences on host infectivity (Pande et al., 2013; Tao et al., 2014). The model human fungal pathogen *Cryptococcus neoformans* grows primarily as a budding yeast, with a filamentous form generally associated with mating and not virulence (Wang et al., 2012). Here we use the seemingly subtle differences between different yeast forms of *C. neoformans* to address the role that phenotypic heterogeneity plays in intra-host dissemination.

A ubiquitous saprophytic fungus associated with trees and bird guano (Emmons, 1955; Lazera et al., 1996), *C. neoformans* is generally harmless, with human exposure thought to be as high as 70% by age five (Goldman et al., 2001). Infection follows inhalation of a fungal cell or spore (Maziarz and Perfect, 2016), and in immunocompromised individuals *C. neoformans* disseminates via the bloodstream and/or lymphatics to almost any organ in the body (Lin et al., 2015; Maziarz and Perfect, 2016). *C. neoformans* cells rapidly proliferate in the central nervous system, and the resulting fungal meningoencephalitis drives cryptococcosis patient mortality, killing ∼180,000 people each year worldwide (Rajasingham et al., 2017).

*C. neoformans* is an excellent model organism to study the impact of phenotypic heterogeneity on intra-host dissemination. Two of its primary virulence attributes—cell and capsule size—vary considerably during infection and are readily observable via microscopy and flow cytometry (Li and Nielsen, 2017). Isolates belonging to the *Cryptococcus* genus exhibit a high degree of cell and capsule size diversity, and morphological heterogeneity strongly impacts *C. neoformans* virulence: increased capacity for morphological variation among isolates correlates with worse clinical outcome (Fernandes et al., 2018).

Cell and capsule size are finely tuned to a multitude of environmental signals and impact stress resistance, immune recognition, and overall pathogenesis (Casadevall et al., 2019; O’Meara and Alspaugh, 2012). Cells grown in nutrient replete laboratory media are morphologically homogeneous (5–10 μm in total (cell + capsule) diameter) and poorly encapsulated (O’Meara and Alspaugh, 2012), while fungal cells proliferating in the lungs are morphologically heterogeneous (1-100 μm in total diameter) and encapsulated, although capsule thickness varies considerably (Wang and Lin, 2015; Zaragoza, 2011). In the host, the polysaccharide capsule shields immunostimulatory microbe-associated molecular patterns in the fungal cell wall, suppresses phagocytosis, and protects against oxidative stress (Casadevall et al., 2019; O’Meara and Alspaugh, 2012).

The best understood *C. neoformans* morphotype is the polyploid (≥4C) “titan” cell, measuring 10-100 μm in cell body diameter (Dambuza et al., 2018; Hommel et al., 2018; Okagaki and Nielsen, 2012; Okagaki et al., 2011; Trevijano-Contador et al., 2018; Zaragoza et al., 2010; Zhou and Ballou, 2018). Titans are more resistant to phagocytosis and oxidative stress and skew immunity toward a non-protective Th2 response (Crabtree et al., 2012; Okagaki and Nielsen, 2012; Okagaki et al., 2010).

While titans contribute considerably to cryptococcal pathogenesis, they are not the only morphotype present during infection, accounting for a minority of cryptococcal cells within the lungs (Zhou and Ballou, 2018). Tissue-specific differences in cryptococcal morphology also manifest in human patients and animal infection models (Denham et al., 2018; Rivera et al., 1998; Xie et al., 2012). Titan cells are rarely observed outside of the lungs, and overall cell and capsule sizes tend to be substantially smaller in extrapulmonary sites, including the brain (Charlier et al., 2005; Denham et al., 2018; Rivera et al., 1998; Xie et al., 2012). While the *C. neoformans* populations in extrapulmonary organs appear to become heterogeneous over time (Lee et al., 1996), the *C. neoformans* population that appears early in the mouse model dissemination process is striking in its homogeneity and small median size (Denham et al., 2018; Fernandes et al., 2018; Fernandes and Carter, 2020; Fernandes et al., 2016; Zaragoza, 2011). We therefore hypothesize that smaller morphotypes are more likely to disseminate and enter extrapulmonary organs.

## RESULTS

### Small cell formation correlates with extrapulmonary dissemination and varies with strain and *Cryptococcus* species

*Cryptococcus* spp. isolates vary in their capacity for morphological variation (Fernandes et al., 2018). To more broadly assess morphological shifts during infection, we inoculated ∼8-week-old female B6 mice with a panel of common *Cryptococcus* spp. reference strains (2.5x10^4^ cells / mouse). At 3 and 17 days post-inoculation (dpi), we homogenized lungs and measured fungal cell body and capsule size (**Figures 1A and 1B**). Morphogenesis varied widely among *Cryptococcus* isolates. While KN99 (*C. neoformans*) and Bt63 (*C. neoformans*) cells shifted smaller between 3 and 17 dpi, R265 (*C. deuterogattii*) and WM276 (*C. gattii*) continued to increase in size—largely due to increased capsule thickness. Smaller median cell size at 17 dpi correlated with increased extrapulmonary burden (**Figures 1C and 1D**). These data suggest that among *Cryptococcus* isolates, the emergence of small cells (<10 μm) within the lungs is a strong predictor of extrapulmonary dissemination.

**Figure 1:**
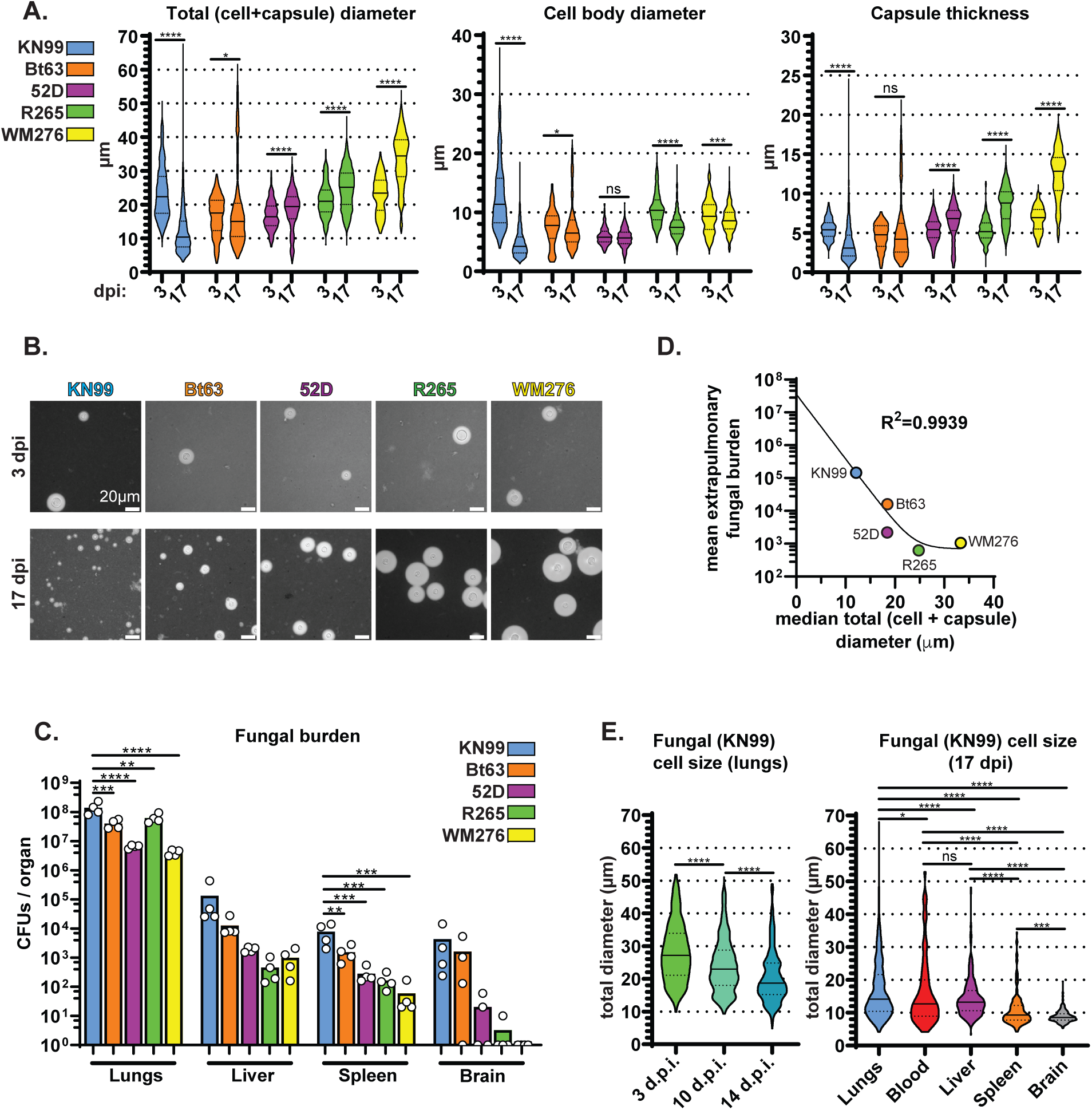
Morphological transitions during pulmonary infections correlate with fungal dissemination. **(A)** Total (cell+capsule) diameter, cell body diameter, and capsule thickness measurements at 3 and 17 days post-intranasal inoculation (dpi) of mice (Mann-Whitney U test; N=4 mice per timepoint, 57-65 fungal cells measured per mouse at 3 dpi and 117-266 fungal cells per mouse at 17 dpi). Solid lines in the violin plots indicate the median and dotted lines mark quartiles. **(B)**Representative India ink images of fungal cells quantified in 1A. **(C)**Fungal burden in mice at 17 dpi (One-way ANOVA and uncorrected Fisher’s LSD). The bar graph represents the mean. **(D)** Nonlinear regression of mean extrapulmonary (liver + spleen + brain) fungal burden and median total (cell + capsule) diameter. **(E)** Total diameter in KN99-infected lungs at 3, 10, and 14 dpi and total diameter in KN99-infected organs at 17 dpi (Mann-Whitney U-test; N=4 mice per timepoint; fungal cells measured per mouse in the lungs: 275, blood: 38, liver: 72, spleen: 43, brain: 72). Solid lines in the violin plots indicate the median and dotted lines mark quartiles. All panels in Figure 1: ns: not significant *p<0.05; **p<0.01; ***p<0.001; ****p<0.0001.

We also measured cell and capsule size in the blood, liver, spleen, and brains of mice inoculated with KN99 cells (the most widely used reference strain) (**Figure 1E**). The range of cell size in the blood and liver was surprisingly broad considering that murine microcapillaries can be as small as 3.5 μm. Although rare, we did find fungal cells in the blood measuring 50 μm across. However, median total diameter in the blood at 17 dpi was 12.72 μm, which was noticeably smaller than earlier timepoints in the lungs (27.21 μm at 3 dpi and 18.71 μm at 14 dpi) indicating that entry into the blood may still be a size-limiting bottleneck. In contrast, cell size in the spleen and brain was far more restricted. The vast majority of cells in the spleen were <15 μm, while cells in the brain rarely exceeded 10 μm in total diameter. These data suggest that the ability to form small cells/capsules in the lungs is a strong predictor of extrapulmonary dissemination, and that multiple bottlenecks may restrict vascular dissemination.

### Inoculum size affects the rate of morphological shift

Lower initial cell density increases the frequency of titan cell formation *in vitro* (Trevijano-Contador et al., 2018) and *in vivo* (Okagaki et al., 2010). We hypothesized that reducing the number of cells in the inoculum would increase the frequency of larger cells early in lung infection and slow the rate of small cell formation (Figures S1A, S1B**, and** S1C). We inoculated mice with 10^3^, 10^4^, or 10^5^ KN99-mCherry cells per mouse and employed the flow cytometry forward scatter-area (FSC-A) parameter as a high-throughput assay to estimate relative changes in cell size at 3, 7, 10, 14, and 17 dpi.

Consistent with our hypothesis, inoculating mice with fewer fungal cells increased fungal cell size at 3 dpi, decreased the rate of cell size reduction in the lungs (Figures S1A, S1B**, and** S1C), decreased lung fungal burden (Figure S1D), and decreased dissemination to the brain at 17 dpi (Figure S1E). Therefore, lower initial fungal cell density slows the rate at which fungal cells shift smaller in the lungs and disseminate to the brain.

### Populations of cells isolated from the lungs display size-dependent differences in cell surface features and differentially bind host factors

When we compared the labor-intensive microscopy-based measurements of cell size with flow cytometry-based measurements, we still observed a fungal size shift over the course of infection (Figure S1A). Thus, we sought to isolate KN99-mCherry cells from infected lung tissue (14 dpi) via fluorescence-activated cell sorting (FACS) using forward scatter area (FSC-A) as a stand-in for size. We sorted cross-sections of the fungal cell size distribution corresponding to the smallest (small *ex vivo*: median total diameter of 7.6 μm), intermediate (mid *ex vivo*: median total diameter of 14.0 μm), and largest (large *ex vivo*: median total diameter of 21.8 μm) ∼20% of the population (**Figure 2A**). We further confirmed that we isolated distinct populations based on cell and capsule size (**Figure 2B**). Fungal cells sorted from each gate were also equivalently viable (**Figure 2C**).

**Figure 2:**
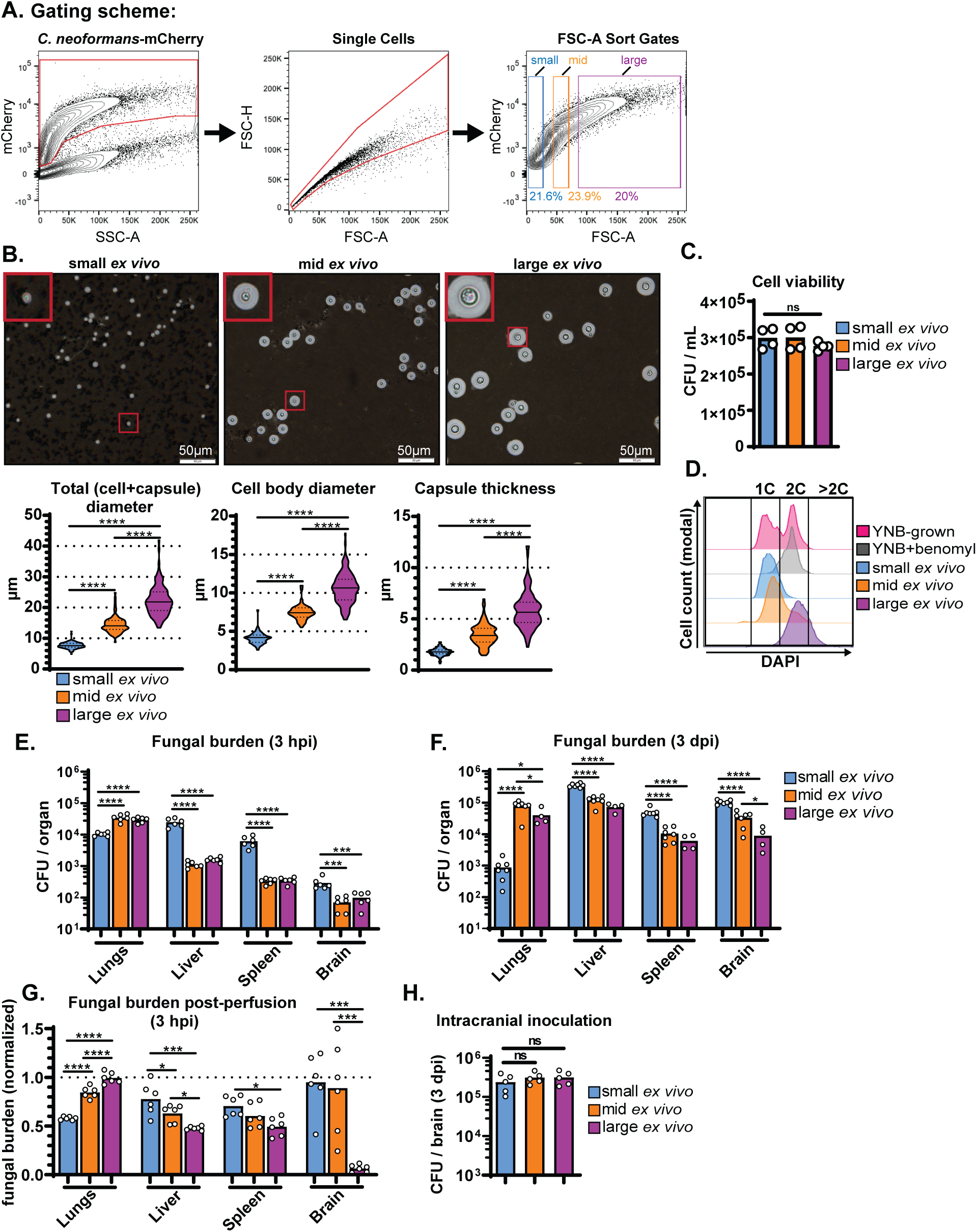
Sorting and characterization of fungal populations from infected lungs. **(A)** FACS gating scheme for sorting KN99-mCherry cells from infected mouse lungs into populations of “small”, “mid”, and “large” cells using forward scatter-area (FSC-A) to approximate size. **(B)** Representative India ink images of cells sorted from “small”, “mid”, and “large FSC-A gates and cell/capsule size measurements (N=97-120 fungal cells). Solid lines in the violin plots indicate the median and dotted lines mark quartiles. **(C)**Cell viability post-sort quantified by colony-forming unit (CFU) count (N=4). **(D)**Representative flow plot of DNA-content measured with DAPI staining. YNB (minimal YNB medium). **(E)** Fungal burden, measured by colony forming unit counts (CFU), at 3 hours post-inoculation (hpi) and 3 days **(F)** post-inoculation (dpi) in mice intravenously-inoculated with small, mid, or large *ex vivo* cells (One-way ANOVA and uncorrected Fisher’s LSD). **(G)** Fungal burden in mice inoculated with sorted *ex vivo* fungal cells and perfused at 3 hpi. Fungal burden was normalized to inoculated, but non-perfused control mice (Mann-Whitney U-test). **(H)** Fungal burden at 3 dpi in mice intracranially inoculated with sorted *ex vivo* fungal cells (One-way ANOVA and uncorrected Fisher’s LSD).

We then measured DNA ploidy with 4′,6-diamidino-2-phenylindole (DAPI) staining (**Figure 2D**). We used haploid cells cultured in minimal yeast nitrogen base (YNB) medium +/-benomyl as a control, since benomyl inhibits cell division and traps cells at 2C (Jacobs et al., 1988). Cells with >2C content are considered titan cells. Only the large *ex vivo* population contained an appreciable number of titan cells (mean: 21.4% >2C) (**Figure S2**). Intermediate *ex vivo* cells averaged 74.2% 1C and 22.8% 2C, while small *ex vivo* cells averaged 89.2% 1C and 10.0% 2C.

### Small *ex vivo* cells disseminate to extrapulmonary organs at a higher rate than intermediate and large *ex vivo* cells

Most infection models—including intravenous dissemination models—use *C. neoformans* yeast cells collected from nutrient replete laboratory media as the inoculum (Mukaremera et al., 2019; Ngamskulrungroj et al., 2012; Sabiiti et al., 2012), failing to capture the phenotypic heterogeneity generated during lung infection. Here we accounted for phenotypic heterogeneity generated during lung infection by isolating cells directly from infected lung tissue and sorting them according to size. To directly address the impact of fungal cell and capsule size on dissemination, we intravenously inoculated naïve mice with small, intermediate, and large *ex vivo* cells (10^5^ cells / mouse). With blood flow from the tail vein running directly to the heart and then lungs for reoxygenation, the lungs harbor the first major bed of microcapillaries fungal cells are likely to encounter following tail vein inoculation. Therefore, sorting fungal cells from infected lung tissue prior to tail vein inoculation roughly replicates the fate of fungal cells had they entered the bloodstream from the lungs. Consistent with our hypothesis, a greater number of small *ex vivo* cells reached the liver, spleen, and brain by 3 hours post-inoculation (hpi) than intermediate and large *ex vivo* cells (**Figure 2E**). In contrast, the majority of intermediate and large *ex vivo* cells remained in the lungs at 3 hpi, possibly because they failed to navigate the lung microcapillaries as well as small *ex vivo* cells. We observed the same trends in fungal burden at 3 dpi, indicating that small *ex vivo* cells are well-capable of proliferating in extrapulmonary tissues (**Figure 2F**).

Although brain fungal burden was equivalent in mice inoculated with intermediate and large *ex vivo* cells at 3 hpi, there were more mid *ex vivo* cells in the brain by 3 dpi. We asked whether the reduced brain fungal burden in mice inoculated with large *ex vivo* cells was due to inefficient blood-brain barrier crossing or reduced proliferation in the brain. To estimate blood-brain barrier crossing *in vivo*, we intracardially perfused mice at 3 hpi to drive out circulating fungal cells and normalized the fungal burden in perfused mice to nonperfused mice inoculated on the same day. While perfusion did not significantly affect brain fungal burden in mice inoculated with small or mid *ex vivo* cells, perfusion drove nearly all of the large *ex vivo* cells from the brain (**Figure 2G**). In contrast, fungal burden post-intracranial inoculation was independent of initial fungal cell size (**Figure 2H**), suggesting that while large *ex vivo* cells are poorer at reaching the brain and crossing the blood-brain barrier, they are not deficient at proliferating in the brain.

### Cell surface differences between *ex vivo* populations result in uptake of small *ex vivo* cells by macrophages

Morphological changes in *C. neoformans* are often accompanied by alterations in cell surface architecture and immune recognition. We screened the binding rate (% total fungal cells bound) of a panel of soluble receptors to small, intermediate, and large *ex vivo* cells (**Figure 3A**). Large *ex vivo* cells were more likely to be bound by complement protein C3 than intermediate, or small *ex vivo* cells (p < 0.05). In contrast, small and intermediate *ex vivo* cells were more likely to be bound by lung surfactant protein D (SPD) (p < 0.01). SPD binding to *C. neoformans* is deleterious for the host, potentially because SPD agglutinates fungal cells and disrupts fungal clearance by macrophages (Geunes-Boyer et al., 2012). We saw low-level, uniform binding of IgA and IgM class antibodies, and little to no binding by mannose-binding lectins.

**Figure 3:**
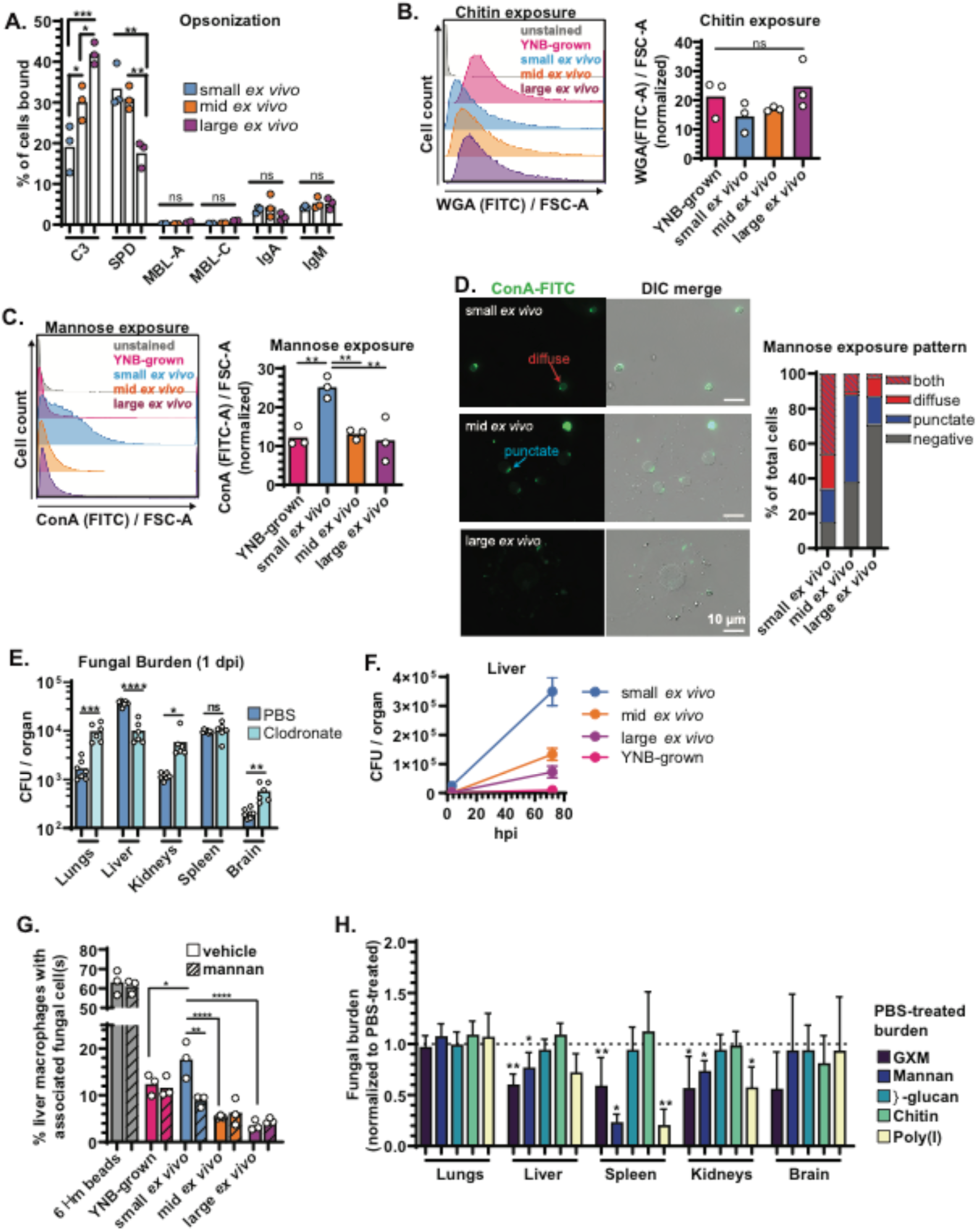
Small *ex vivo* cells disseminate to extrapulmonary organs at a higher rate than larger cells, with mannose recognition by macrophages contributing to their clearance from the vasculature. **(A)** Percentage of total cells bound by complement (C3), surfactant protein D (SPD), mannose binding lectins A and C (MBL-A; MBL-C), IgA, and IgM. **(B)** Representative flow plots and quantification of wheat-germ agglutinin (WGA) stained cells to determine chitin exposure. **(C)**Representative flow plots and quantification of concanavalin A (ConA) stained cells to determine mannose exposure. **(D)**Representative images and characterization of mannose exposure pattern after staining with ConA (N=50 fungal cells). All comparisons in Figure 2 were analyzed using one-way ANOVA and uncorrected Fisher’s LSD (ns: not significant; *p<0.05; **p<0.01; ***p<0.001; ****p<0.0001). All bar graphs display the mean. **(E)** Fungal burden at 1 dpi in mice pretreated with either clodronate or vehicle control (PBS) liposomes and then inoculated with small *ex vivo* cells and (Unpaired t test). **(F)** Fungal burden in the livers of mice (N=4-6) inoculated with *ex vivo* fungal cells or fungal cells harvested from minimal YNB (YNB) medium at 3 and 72 hpi. Points indicate the mean and the error bars indicate standard deviation. **(G)** Percentage of primary cultured murine liver macrophages with physically associated plastic beads or fungal cells (stained with Direct Yellow 96) after 4 hours. Dashed lines indicate 10-minute pretreatment with exogenous mannan (One-way ANOVA and uncorrected Fisher’s LSD; N=2263-3197 total macrophages counted across 3 independent replicates). **(H)** Fungal burden in mice intravenously treated with indicated ligand 2-3 minutes prior to intravenous inoculation with small *ex vivo* fungal cells. Mice were perfused 10 minutes post-inoculation and fungal burden was normalized to vehicle (PBS) treated controls (Unpaired t-test; N=6 mice per group). All panels: ns: not significant *p<0.05; **p<0.01; ***p<0.001; ****p<0.0001. All bar graphs display the mean.

Differences in host factor binding suggested that cell surface architecture may differ among the sorted populations. We used two lectins to probe the exposure of microbial features in different layers of the cell wall: wheat germ agglutinin (WGA), which recognizes chitin in the innermost layers of the cryptococcal cell wall and concanavalin A, which binds mannose on mannoproteins in the outer cell wall and inner capsule. There were no significant differences in chitin exposure as measured by flow cytometry (**Figure 3B**). However, mannose was significantly more exposed on small *ex vivo* cells compared with intermediate and large *ex vivo* cells and cells grown in non-capsule inducing minimal YNB medium (**Figure 3C**).

We visualized mannose exposure patterns via microscopy and characterized staining as negative, punctate (one or more distinct staining puncta), diffuse (staining covers the observable cell surface), or both punctate and diffuse (**Figure 3D**). Mannose was more likely to be exposed both at specific puncta and diffusely across the surface of small *ex vivo* cells. Mannose exposure can correspond to recent bud scars, which may explain the punctate staining in our analysis (Panepinto et al., 2007). The high prevalence of diffuse mannose exposure on small cells may be due to their relatively thin capsules. Altogether, these results suggest that cell and capsule size heterogeneity correlates with heterogeneity in cell surface architecture and immune recognition.

As small *ex vivo* cells most efficiently disseminated to extrapulmonary organs, we examined the mechanisms governing their dissemination profile more closely. Most small *ex vivo* cells that made it past the lungs at 3 hpi were cleared in the liver and spleen (**Figure 3E**). The liver plays a major role in clearing microbes and microbial components from the blood (Macpherson et al., 2016) via recognition of microbial-associated molecular patterns (MAMPs) (Kubes and Jenne, 2018). Recently, intravital imaging revealed that liver macrophages clear *C. neoformans* cells from the bloodstream (Sun et al., 2019). However, the inoculum for those studies consisted of *C. neoformans* cells cultured *in vitro*. Our cell surface characterization revealed that small *ex vivo* cells differ from cells cultured *in vitro* (YNB medium), intermediate *ex vivo*, and large *ex vivo* cells, which may influence host cell recognition of *C. neoformans* cells.

Phagocytes such as platelets, neutrophils, and macrophages are critical for the clearance of various pathogens. To identify those important for small cell clearance, we systematically depleted each cell type and measured organ distribution of *ex vivo* populations following tail vein injection. To address the role of macrophages, we depleted macrophages from the liver, spleen, and bone marrow of mice with clodronate liposomes 48 hours prior to inoculation (**Figures S3A and S3B**). Macrophage depletion reduced small *ex vivo* cell fungal burden in the liver, while increasing fungal burden in the lungs, kidneys, and brain at 1 dpi (**Figure 3E**). These results corroborate the findings of Sun *et al*, in that liver macrophages play a critical role in filtering *C. neoformans* from the blood (Sun et al., 2019). However, we also found that small *ex vivo* cells grow rapidly in the liver compared with *in vitro*-grown cells or other *ex vivo* populations (**Figure 3F**), suggesting that attempts by the host to clear *C. neoformans* cells by sequestering them in the liver may backfire.

Platelets can recognize either complement protein bound to microbes or microbial features directly to initiate microbial clearance, especially in the spleen (Broadley et al., 2016). Platelets don’t appear to interact with encapsulated *C. neoformans*, or mediate the clearance of *in vitro*-grown *C. neoformans* cells from the blood (Sun et al., 2019). When we depleted platelets, we also did not see any effect on fungal dissemination (**Figures S4A-C**).

It is unclear whether neutrophils play a predominantly protective or detrimental role during cryptococcosis, though there is evidence neutrophils can clear *C. neoformans* from the blood (Sun et al., 2016; Zhang et al., 2016). Neutrophil depletion greatly increased fungal burden in the lungs and brain when *in vitro*-grown fungal cells were injected intravenously (Sun et al., 2016; Zhang et al., 2016). However, neutrophil-depletion prior to inoculation with small *ex vivo* cells slightly increased lung fungal burden but did not affect fungal burden in other organs, indicating that the small *ex vivo* cells are more resistant to neutrophil-mediated clearance than typical *in vitro*-grown fungal cells (**Figures S4D-F**).

### Macrophage recognition of small *ex vivo* cells is partially mediated by mannose exposure

We predicted that liver macrophages may recognize the high levels of exposed mannose on the surface of small *ex vivo* cells. We isolated liver macrophages from healthy ∼8-week-old B6 mice (Li et al., 2014) and co-cultured them with 6 μm plastic beads, YNB-grown cells, and sorted *ex vivo* populations at a multiplicity of infection of 2:1. Half of the wells also received pretreatment with exogenous mannan, with the prediction that it would reduce macrophage association if mannan recognition is critical. After 4 hours we washed away free fungal cells, leaving only the fungal cells bound or internalized by macrophages, which we define as “macrophage association”.

Approximately 60% of macrophages had associated with 6 μm plastic beads by 4 hours, indicating that the liver macrophages still displayed phagocytic capacity *ex vivo* (**Figure 3G**). Additionally, 18% of liver macrophages associated with small *ex vivo* cells, which was more than twice the frequency of intermediate and large *ex vivo* cells. Pretreatment with exogenous mannan prior to infection reduced macrophage association with small *ex vivo* cells by ∼40%, but did not affect association with intermediate and large *ex vivo* cells, YNB-grown cells, or beads. These data suggest that liver macrophages recognize small *ex vivo* cells in part through exposed mannose, and that mannose exposure could contribute to fungal sequestration in the liver.

Liver macrophages and other macrophages that sample the bloodstream are subject to shear force, which can affect recognition and phagocytosis of microbes and other foreign matter (Broadley et al., 2016). We assessed whether mannose exposure on small *ex vivo* cells affects vascular clearance *in vivo* by intravenously injecting either 400 μg of exogenous mannan or vehicle (PBS) 2-3 minutes prior to intravenous inoculation with 10^5^ small *ex vivo* cells. 10 minutes later, we perfused mice to drive out non-recirculating fungal cells and plated organs for colony forming units (CFUs) to estimate fungal burden. If mannose recognition contributed to fungal clearance in a given organ, we expected to see reduced fungal burden in mice that received exogenous mannan, compared with the vehicle control. We repeated this with different fungal cell wall and capsule components (GXM, β-glucan, chitin), as well as polyinosinic acid (poly(I)), which is a scavenger receptor ligand (Pearson et al., 1993).

Although mannan did reduce fungal clearance in the liver and kidneys by ∼20% and in the spleen by ∼75% (**Figure 3H**), it was not the only fungal ligand that influenced vascular clearance. Poly(I) reduced clearance to a strikingly similar extent as mannan. The purified capsule polysaccharide GXM reduced clearance in the liver, spleen, kidneys at roughly the same level (∼40% reduction). β-glucan and chitin, on the other hand, did not appreciably affect fungal clearance, possibly because of their buried position beneath the capsule. Fungal burden in the brain was too low – usually less than 100 CFU per mouse -to detect changes. These data indicate that clearance of small *ex vivo* cells is not dependent on a single cell surface ligand and that there are tissue-specific dependencies on various fungal features that mediate vascular clearance.

### Enhanced small *ex vivo* cell dissemination is not solely dependent on size

We next assessed whether small *ex vivo* cells disseminate more efficiently due to their size alone, or if there were other contributing factors. We inoculated mice via the tail vein with either large *ex vivo* cells, small *ex vivo* cells or size-matched inert polystyrene beads. Large *ex vivo* cells disseminated similarly to size-matched 25 μm beads; the majority resided in the lungs at 3 hpi (**Figure 4A**). Additionally, more 6 μm beads traveled past the lungs to reach the liver and spleen compared to 25 μm beads. However, small *ex vivo* cells reached the liver and spleen at an even higher frequency, indicating that absolute size influences fungal dissemination through the vasculature, but it is not the only contributing factor.

**Figure 4:**
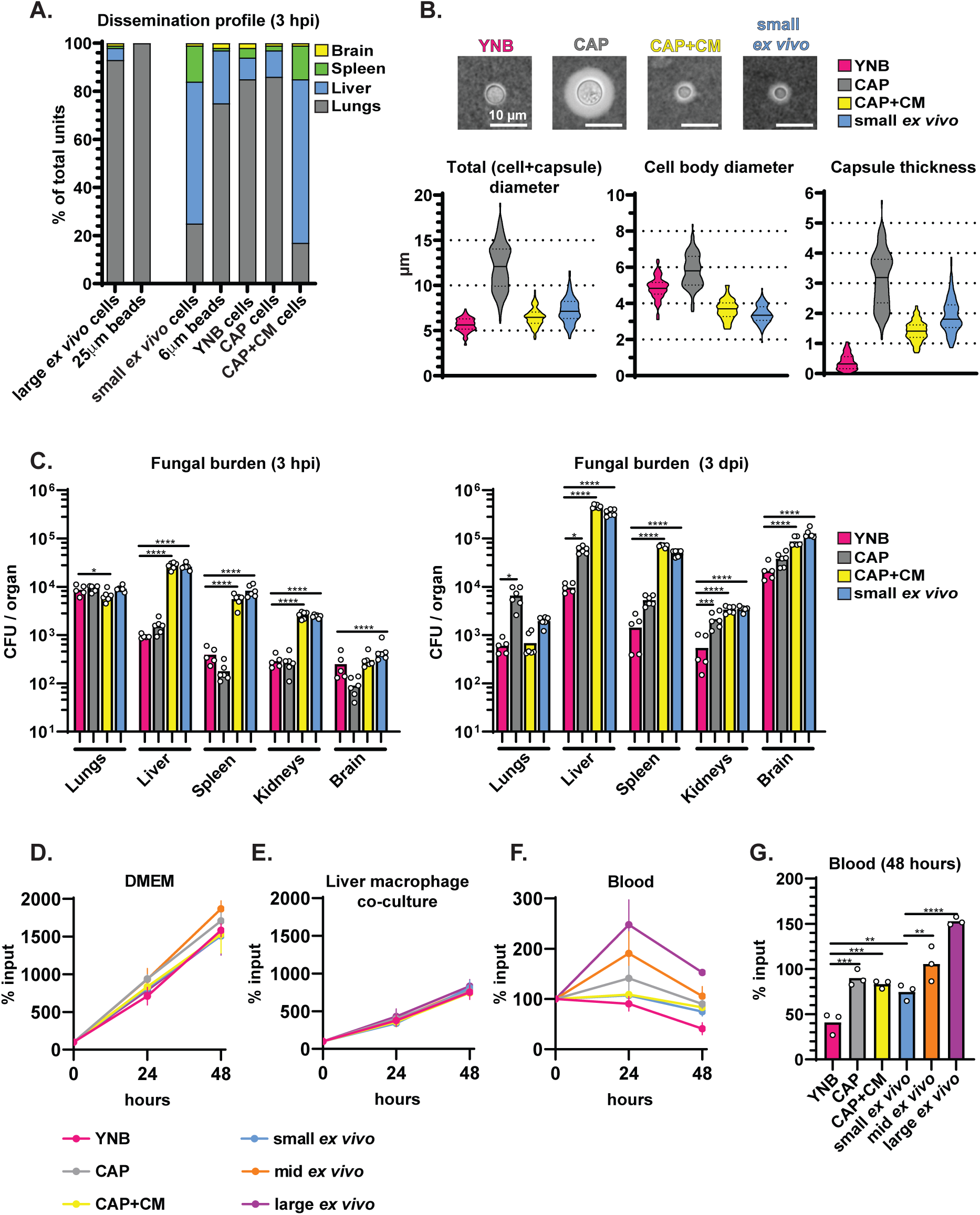
Small cells generated under host-like conditions *in vitro* replicate the dissemination profile of small *ex vivo* cells. **(A)** Percentage of total plastic beads or live cells in the indicated organ 3 hours post-intravenous inoculation (hpi). **(B)**Representative India ink images and quantification of cell and capsule size (N=100 fungal cells). YNB (minimal YNB medium); CAP (capsule inducing medium); CAP+CM (capsule inducing medium supplemented with 10% conditioned medium). **(C)** Fungal burden, measured by colony forming units (CFU), in mice at 3 hpi. and 3 days post-inoculation (dpi). **(D)**Fungal cell survival in Dulbecco’s modified Eagle’s medium (DMEM) and **(E)** liver macrophage co-culture (multiplicity of infection of 1:1) measured as a percentage of input. **(F)** Fungal cell survival in mouse blood measured as a percentage of input over 48 hours and **(G)** at 48 hours growth. All comparisons in Figure 4 were analyzed using one-way ANOVA and uncorrected Fisher’s LSD (*p<0.05; **p<0.01; ***p<0.001; ****p<0.0001). All bar graphs display the mean.

We next examined *in vitro* conditions that could reproduce the small *ex vivo* cell morphotype and dissemination profile. Previously, we observed that cells grown in capsule-inducing medium (CAP; 10% Sabouraud’s dextrose broth buffered to pH 7.4) shifted toward smaller cell and capsule size if they were supplemented with conditioned medium (CM) from cells grown in non-capsule inducing medium (YNB) after 24 hours (Denham et al., 2018). YNB medium contains vitamins, nitrogen, and glucose but not amino acids. YNB-grown cells suppress capsule formation, and are typically <10 μm in diameter. CAP medium induces capsule formation and is sufficient for cell division. We collected CM by filtering the supernatant from saturated YNB-grown cultures. In this case, CM consists of unspent nutrients and soluble fungal factors secreted during growth in non-capsule inducing conditions.

We compared the dissemination profile of small *ex vivo* cells to cells grown in three *in vitro* populations: (1) YNB-grown cells (2) CAP-grown cells, and (3) CAP-grown cells supplemented with CM (CAP+CM). We also subjected the *in vitro*-grown populations to the same sorting process as *ex vivo* cells by sorting YNB, and CAP+CM-grown cells from the “small” FSC-A gate (YNB cells: 5.6 μm median total diameter; CAP+CM cells: 6.5 μm median total diameter; few YNB or CAP+CM cells fell in the “mid” or “large” FSC-A gates) and sorting the CAP-grown cells from the “mid” FSC-A gate (12.1 μm median total diameter; few CAP-grown cells fell into the “small” or “large” FSC-A gates) (**Figure 4B**).

Culture conditions and morphology of cells in the inoculum heavily impacted dissemination. Only CAP+CM cells replicated the dissemination profile of small *ex vivo* cells in terms of fungal burden at 3 hours and 3 days post-inoculation (**Figure 4C**). Despite being roughly the same size as small *ex vivo* cells, YNB-grown cells displayed similar fungal burden to CAP-grown cells at 3 hours, and lower fungal burden in extrapulmonary organs than CAP, CAP+CM, and small *ex vivo* cells at 3 days. Therefore, both absolute fungal cell size, and the physiological state of cells entering the blood influence the outcome of vascular dissemination.

### Host-adapted cells survive better in the blood than cells grown in nutrient-replete medium

The caveat to using fungal burden as a readout for dissemination is that we cannot easily distinguish between the ability of cells to reach certain organs and their ability to survive in those organs. We hypothesized that YNB-grown cells may not be as primed for survival in the bloodstream as cells sorted from infected host-tissue or cells grown under capsule-inducing medium (CAP medium).

To address that hypothesis, we examined the survival of *in vitro-* and *ex vivo*-sorted cells in whole mouse blood or liver macrophage co-culture (multiplicity of infection of 1:1). We used DMEM as a tissue culture medium control (96-well plates, 37 °C, 5% CO_2_, static). We assessed fungal cell survival as a percentage of input at 0-, 24-, and 48-hour timepoints. Liver macrophages suppressed *in vitro* and *ex vivo* cell growth equivalently (**Figures 4D and 4E**).

However, the *ex vivo* populations and *in vitro* populations from host-like conditions survived better in blood than YNB-grown cells (**Figure 4F and 4G**), with large and intermediate *ex vivo* populations faring the best. These results support the idea that size and physiology are critical for bloodstream dissemination: small *ex vivo* cells potentially combine the stress resistance required for increased survival in the blood with the smaller cell size required for widespread dissemination via microcapillaries.

### Expression of phosphate acquisition genes differ between small cells and larger *C. neoformans* cells

Since CAP+CM-grown cells resemble small *ex vivo* cells in terms of cell + capsule size and vascular dissemination, we hypothesized that they might suitably model small cell formation in the lungs. We analyzed the transcriptome of cells grown in YNB, CAP, and CAP+CM medium to identify genes that might facilitate the transition to small cell size in host-like capsule inducing conditions. We also analyzed the transcriptome of small, intermediate and large *ex vivo* cells.

Comparing the transcriptome of CAP, CAP+CM, and the *ex vivo* populations to YNB-grown cells demonstrated that the YNB-grown transcriptome is vastly different from each of these populations (**Figure 5A** and **5B**). There were also 1085 up-regulated and 1297 downregulated genes shared only among the *ex vivo* populations relative to YNB-grown cells. Core sets of 699 and 291 genes were up- and down-regulated respectively among the *ex vivo* populations and CAP/CAP+CM populations relative to YNB-grown cells, demonstrating that the transcriptome within the host environment is quite distinct from *in vitro* conditions that induce virulence traits. Among the *ex vivo* populations, small *ex vivo* cells were the most transcriptionally distinct (**Figure 5A**). 928 genes were differentially expressed in small vs intermediate *ex vivo* cells, and 2,365 genes were differentially expressed in small vs large *ex vivo* cells. For comparison, only 93 genes were differentially expressed in intermediate vs large *ex vivo* cells (**Figure 5A**).

**Figure 5:**
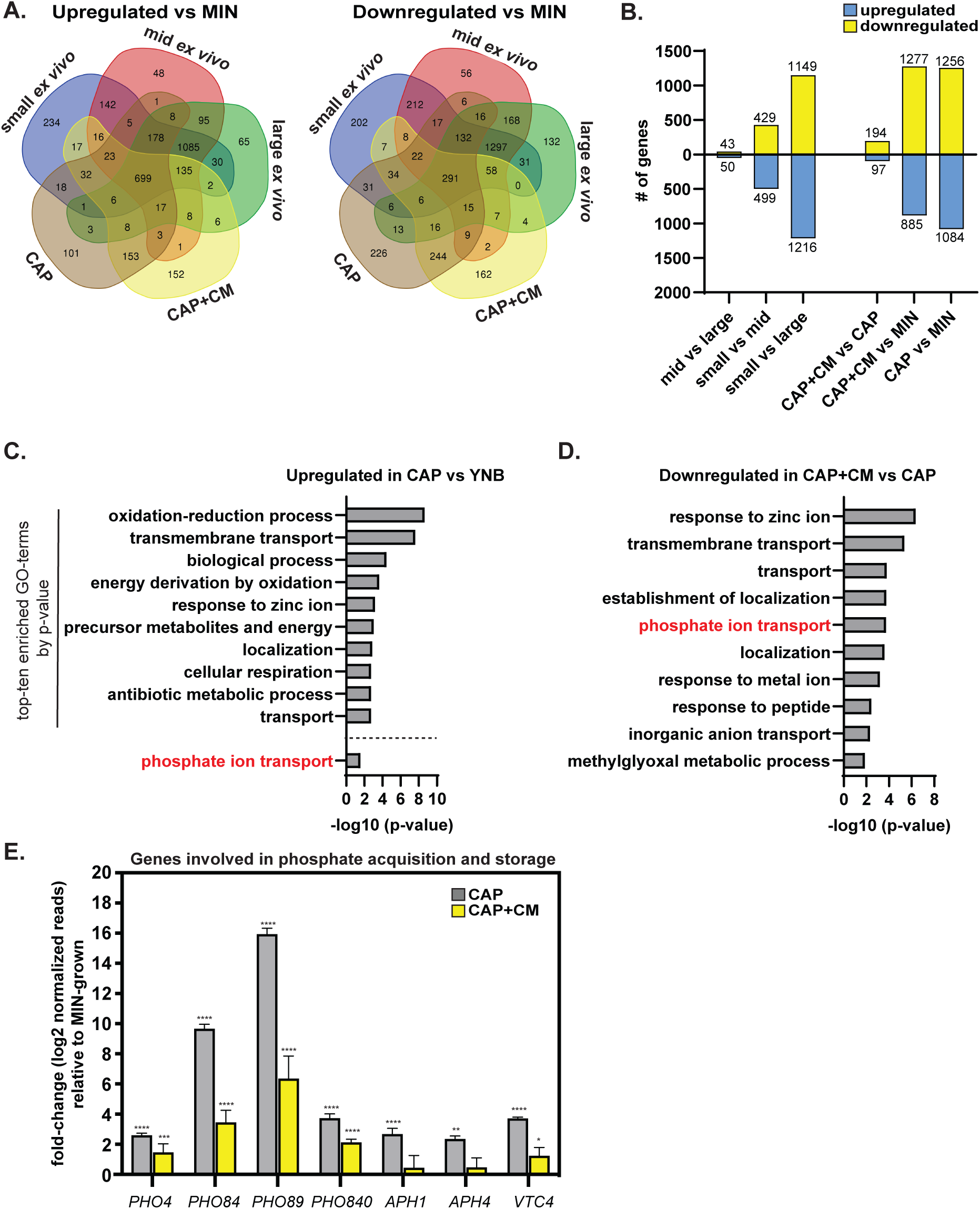
Expression of phosphate acquisition genes correlate with cell and capsule size changes *in vitro* and are upregulated *in vivo*. **(A)** Venn diagram of genes upregulated or downregulated in different conditions relative to YNB-grown cells (YNB: minimal YNB medium; CAP: capsule-inducing medium; CAP+CM: capsule inducing medium supplemented with 10% conditioned medium). **(B)** Comparisons of the total number of differentially regulated genes among *in vitro*-grown populations. **(C)** Top-ten most significantly enriched gene ontology (GO) terms representing genes upregulated in CAP vs YNB-grown cells (-log10 (p-value)). Phosphate ion transport was #46. **(D)**Top-ten most significantly enriched GO-terms representing genes downregulated in CAP+CM vs CAP-grown cells (-log10 (p-value)). **(E)** Select phosphate acquisition and storage RNA-seq gene expression relative to YNB-grown cells (adjusted p values; *p<0.05; **p<0.01; ***p<0.001; ****p<0.0001). All bar graphs display the mean and error bars the standard deviation.

Gene ontology term (GO-term) enrichment analysis revealed that phosphate ion transport was upregulated in CAP-grown cells vs YNB-grown cells (**Figure 5C**), and downregulated in CAP+CM vs CAP-grown cells (**Figure 5D**). In other words, phosphate acquisition genes are induced when cells enter capsule-inducing conditions (CAP medium), and then suppressed following supplementation with CM.

We observed the same trends when analyzing the expression of specific genes involved in phosphate acquisition and storage (**Figure 5E**). When *C. neoformans* cells are starved of phosphate, the phosphate acquisition transcription factor, Pho4, translocates to the nucleus where it promotes its own gene expression and the expression of genes involved in phosphate acquisition and storage. These include phosphate transporters: *PHO84*, *PHO89*, and *PHO840*; phosphatases: *APH1* (secreted/vacuolar acid phosphatase) and *APH4* (predicted intracellular acid phosphatase); and *VTC4* (vacuolar transport chaperone involved in processing polyphosphate). Pho4-dependent genes were upregulated in CAP vs YNB-grown cells and then suppressed in following CM supplementation—although many phosphate acquisition genes still remained more highly expressed in CAP+CM-grown cells compared with YNB-grown cells.

Phosphate drives *C. neoformans* populations toward smaller morphotypes

We hypothesized that phosphate might be a critical nutrient that facilitates the emergence of small cells in CAP+CM media. Conditioned medium consists of unspent YNB medium and soluble fungal cell derived factors (Albuquerque et al., 2013). YNB medium is defined, so we screened every component individually (including phosphate) to determine which components are sufficient to induce small cell formation when cells are grown in CAP medium. We also tested two soluble fungal-derived factors that accumulate during growth in YNB medium: exo-GXM (Denham et al., 2018) and the quorum sensing-like peptide Qsp1 (Homer et al., 2016; Lee et al., 2007).

We grew cells for 24 hours in YNB medium, then sub-cultured 10^5^ cells/mL into CAP medium for another 24 hours. We then sub-cultured CAP-grown cells 1:1 into fresh CAP medium with 10% of the final volume being a YNB medium component, fungal-derived factor, or water as the vehicle control. After a final 24-hour period, we measured cell size (FSC-A) by flow cytometry.

Conditioned YNB medium and fresh YNB medium were sufficient to stimulate a shift toward smaller cell size (**Figures 6A and 6B**). Among the YNB medium components, only phosphate was sufficient to stimulate a shift toward smaller cell size (**Figure S5A**), and did so in a concentration-dependent manner (**Figures 6A** and **6B**). Phosphate-supplemented cells decreased both total (cell+capsule) diameter (**Figure 6C**) as well as cell (**Figure S6A**) and capsule (**Figure S6B**) size individually. Phosphate was also sufficient to induce a shift toward smaller size for cells grown in titan cell-inducing medium (5% Sabouraud’s, 10% fetal calf serum, 15 μM sodium azide, pH 7.4, 5% CO2, 37 °C) (**Figure S5B**).

**Figure 6:**
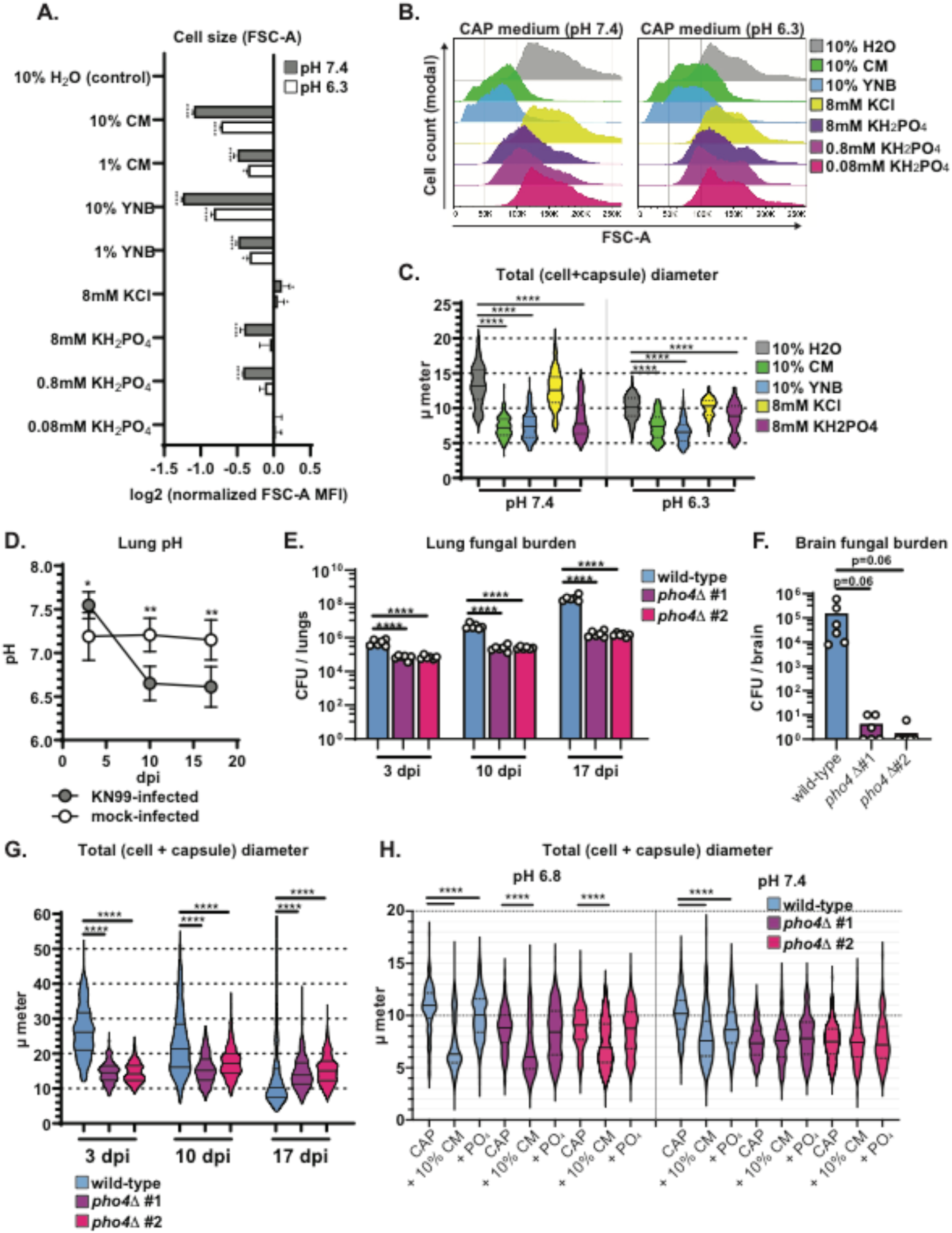
Phosphate is sufficient to drive *C. neoformans* toward smaller morphotypes in alkaline host-like conditions and is required for morphogenesis *in vivo*. **(A)** Cell sizemeasured using flow cytometry forward scatter (FSC-A) after growth in CAP-medium (10% Sabouraud’s pH 7.4) and supplementation with the indicated factor (YNB: minimal YNB medium; CAP: capsule-inducing medium; CAP+CM: capsule inducing medium supplemented with 10% conditioned medium). FSC-A median fluorescence intensity (MFI) was normalized to control (10% H2O)-treated cells (one-way ANOVA and uncorrected Fisher’s LSD). Bar graphs display the mean. **(B)**Representative flow cytometry plots for conditions in Figure 6A. **(C)**Total (cell + capsule) diameter measurements representing conditions in Figure 6A (Mann-Whitney U-test; N=200 fungal cells). Solid lines in the violin plots indicate the median and dotted lines mark quartiles. **(D)** pH in *C. neoformans* strain KN99-infected and mock-infected mouse lungs at 3, 10, and 17 days post-inoculation (dpi) (N=4 mice; one-way ANOVA and uncorrected Fisher’s LSD; error bars represent standard deviation). **(E)** Lung fungal burden in mice at 3, 10, and 17 dpi (One-way ANOVA and uncorrected Fisher’s LSD). The bar graph represents the mean. **(F)** Brain fungal burden in mice at 17 dpi (One-way ANOVA and uncorrected Fisher’s LSD). The bar graph represents the mean. **(G)** Total diameter (cell + capsule) measurements at 3, 10, and 17 dpi of mice (Mann-Whitney U test; N=6 mice per timepoint; 50, 100, and 150 wild-type cells measured per mouse at days 3, 10, and 17 respectively. 50, 50, and 100 *pho4*Δ mutant cells measured per mouse at days 3, 10, and 17 respectively) **(H)** Total (cell + capsule) diameter measurements of wild-type and *pho4*Δ mutant cells after growth in CAP-medium (10% Sabouraud’s at labeled pH, “control”) and supplementation with the indicated factor (10% conditioned medium or 800 μM PO4 at **(H)** pH 7.4 or **(I)** pH 6.8 (Mann-Whitney test; N = 3 biological replicates per experiment and >100 cells measured per replicate. The solid line represents the population median and the dotted lines the 25^th^ and 75^th^ percentiles.). All comparisons marked as significant represent populations for which the 95% confidence interval does not overlap. All panels: ns: not significant *p<0.05; **p<0.01; ***p<0.001; ****p<0.0001.

We previously reported that exo-GXM may trigger the formation of smaller cells (Denham et al., 2018). However, we were unable to reproduce those results (**Figure S5A**), possibly due to differences in exo-GXM preparation, but more likely because we previously solubilized exo-GXM in phosphate buffered saline. Qsp1 is a cell-density dependent signaling peptide that suppresses the formation of titan cells (Trevijano-Contador et al., 2018). However, the addition of Qsp1 did not affect cell size in this model (**Figures S5A and S5B**).

The Pho84 and Pho840 phosphate transporters are phosphate/H^+^ symporters, and *C. neoformans* therefore requires a proton gradient to efficiently transport phosphate into the cell (Kretschmer et al., 2014). In alkaline pH, *C. neoformans* struggles to efficiently import phosphate and upregulates phosphate acquisition machinery (Kretschmer et al., 2014; Lev et al., 2017). *C. neoformans* releases metabolites that acidify the local microenvironment and increase nutrient uptake (Himmelreich et al., 2003; Wright et al., 2002). For instance, cryptococcomas in the brain acidify to levels as low as pH 5.5 (Himmelreich et al., 2003; Wright et al., 2002).

We measured lung pH in mice inoculated with *C. neoformans* throughout infection, and found that pH in the lungs decreased from ∼7.4 to as low as ∼6.3 between 3 and 17 dpi (**Figure 6D**). Phosphate did not induce significant shifts in FSC-A-measured cell size when CAP medium was buffered to 6.3, potentially because phosphate availability in CAP medium buffered to pH 6.3 is not as limiting as pH 7.4 (**Figures 6A and 6B**). However, we did measure a modest, but significant decrease in total (cell+capsule) diameter by light microscopy when pH 6.3 CAP-grown cells were supplemented with phosphate (**Figure 6C**), which was mostly due to a decrease in capsule thickness (**Figure S5**). When phosphate availability suddenly increases due to higher extracellular concentrations, *C. neoformans* cells shift toward smaller morphotypes.

### Limiting phosphate acquisition genetically suppresses morphogenesis *in vivo*

Since phosphate was sufficient to trigger cell and capsule size reduction *in vitro*, we hypothesized that limiting phosphate acquisition *in vivo* would alter morphogenesis, perhaps delaying the appearance of small morphotypes. To test this hypothesis, we intranasally inoculated mice with wild-type KN99, or either of two independently constructed KN99:*pho4*Δ strains, which lack the gene encoding the transcription factor *pho4*. Without *pho4*, *C. neoformans* cells fail to adequately upregulate genes involved in phosphate acquisition when available phosphate is limiting, and become hypersensitive to alkaline conditions. The *pho4*Δ mutant is attenuated for growth in the lungs but more severely attenuated for dissemination to the brain, due to its poor survivability in alkaline blood (Lev et al., 2017). Our findings replicated the work from Lev *et al*., in that the *pho4*Δ mutant cells were attenuated for proliferation in the lungs **(Figure 6E)** and dissemination to the brain **(Figure 6F)**. We also found that cell and capsule size of *pho4*Δ mutant cells populating the lungs was static and less heterogeneous relative to wild-type cells **(Figures 6G**, **S5D-F)**. Most *pho4*Δ mutant cells remained between 10 and 20 μm in total (cell + capsule) diameter over 17 days **(Figure 6G)**, with many of them also exhibiting irregular budding (**Figure S5F**). Total median wild-type cell + capsule diameter was far larger than the *pho4*Δ cells at day 3 (wild-type: 26 μm; *pho4*Δ #1: 15 μm; *pho4*Δ #2: 14 μm) and smaller than the *pho4*Δ cells at day 17 (wild-type: 10 μm; *pho4*Δ #1: 14 μm; *pho4*Δ #2: 15 μm). *pho4*Δ cells therefore appear to be deficient in titan cell formation *in vivo* (**Figure 6G**). However, this result potentially obscures whether *pho4*Δ mutant cells are able to form small cells.

In our *in vitro* small cell induction system, titan cell formation is not required. To eliminate the possibility that *pho4*Δ cells cannot form small cells because they cannot form titan cells, we induced small cells with either conditioned medium or inorganic phosphate. At pH 7.4, *pho4*Δ cells exhibit constitutively small cell bodies and capsules. *pho4*Δ cells do not get substantially smaller in response to phosphate or conditioned medium (**Figure 6H**). At pH 6.8, the upper pH range in the lungs at 10 dpi (**Figure 6D**), we found that *pho4*Δ cells’ total diameters were only slightly smaller than wild-type cells at pH 6.8 (wild-type: 10.6 μm, *pho4*Δ #1: 9 μm; *pho4*Δ #2: 9 μm), while the difference was more dramatic at pH 7.4 (wild-type: 10.2 μm, *pho4*Δ #1: 7.3 μm; *pho4*Δ #2: 7.5 μm) (**Figure 6I**). At pH 6.8, both wild-type and *pho4*Δ cells became smaller in response to conditioned medium but not phosphate, while pH 7.4-grown *pho4*Δ did not exhibit a cell size shift in response to either conditioned medium or phosphate (**Figure 6H**). Together, these data demonstrate that *PHO4* is necessary for phosphate-induced small cell formation at pH 7.4 and that this phosphate-induced pathway could well be the only molecular trigger of seed cell formation at pH 7.4. Our observation that *pho4*Δ cells’ inability to form small cells at pH 7.4 is suppressed at pH 6.8 could indicate that two pathways exist to drive small cell formation. However, since *pho4*Δ cells’ growth defects are suppressed at acidic pH (Lev et al., 2017), Pho4’s transcriptional repertoire and therefore *pho4*Δ cells’ inability to form small cells could also be suppressed.

Finally, we measured phosphate levels within *C. neoformans* cells during growth under capsule-inducing conditions (10% Sabouraud’s pH 7.4 or pH 6.8) and following *in vitro* induction of small cells. We found that small cells contained a higher concentration of phosphate on a per cell basis, despite their smaller cell volume (**Fig. S5G**). Small cell formation therefore involves increased acquisition of a limiting nutrient and is triggered by multiple common signals, strongly supporting the idea that it is a major pathway that can be activated under a variety of conditions.

### Components of *C. neoformans*’s environmental niche induce the small cell morphotype

Phosphate is potentially available from a number of sources that *C. neoformans* could encounter either in the environment or during infection. These include nucleotide pools within both fungal cells and mammalian tissues and cells (Beis and Newsholme, 1975; Harris et al., 1958) and phosphate in the blood at concentrations in the millimolar range (Beis and Newsholme, 1975). Moreover, the infection process itself could increase the amount of extracellular phosphate within host tissues. Neutrophil extracellular traps (NETs) include extruded neutrophil DNA (Brinkmann et al., 2004). Cell lysis releases intracellular nucleoside mono-, di-, or triphosphates, with ATP in particular acting as a signal of damage (Grygorczyk et al., 2021). Extracellular ATP is released by host cells during inflammation (Dosch et al., 2018), in response to injury (Gault et al., 2014), by macrophages in response to bacterial infection (Ren et al., 2014), and as an apoptotic signal (Elliott et al., 2009). Outside the host, *C. neoformans* is often found in association with pigeon nests and guano (Emmons, 1955). Bird guanos are rich in phosphate (Otero et al., 2018). Nucleotide sugars are precursors for critical fungal cell wall components. UDP-GlcNAc is polymerized into chitin (Gow et al., 2017). UDP-glucose is important for β-glucan (Agustinho et al., 2018) and GXM synthesis (Agustinho et al., 2018; Chung and Brown, 2020). However, GXM is synthesized in the secretory compartments (Yoneda and Doering, 2006), so it is possible that exogenous UDP-sugars are not incorporated into GXM or GXMGal because of the challenges of transporting them from the extracellular space to the ER and Golgi (Agustinho et al., 2018). β-glucan and chitin are synthesized at the plasma membrane, so extracellular precursors could be directly incorporated into these structures.

Phosphate availability also varies widely with pH. In the soil environment, phosphate is bound by calcium at alkaline pH and aluminum, iron, and manganese at acidic pHs (Becquer et al., 2014; Hasan et al., 2016). In the host, pH is tightly controlled, but certain conditions, including lung infections, cause respiratory acidosis and decrease the local pH (André et al., 2022). We observe such a pH decrease in the lungs of *C. neoformans*-infected mice (**Fig. 6D**).

Using our *in vitro* small cell induction protocol, we tested whether or not a variety of different phosphate sources could trigger small cell formation. *E. coli* genomic DNA and sheared salmon sperm DNA (ssDNA) were both able to induce a modest but significant shift in the median total diameter of the *C. neoformans* population, from 9.4 microns (at pH 7.4) to 8.0 microns for *E. coli* genomic DNA and 8.3 microns for ssDNA (**Fig. 7A**). Of the nucleoside mono-, di-, and triphosphates, nucleoside triphosphates (NTPs) induced the greatest reduction in median population diameter at both pH 6.8 (8.4 to 7.1 microns) and pH 7.4 (9.4 microns to 6.7 microns) (**Fig. 7B,C**). This appears to be a synergistic effect of multiple NTPs, as the median diameter shift induced by all four NTPs together was greater than any individual NTP (**Fig. S6A,B**). Among the nucleotide sugars, UDP-GlcNAc, a precursor for chitin, induced the largest shift in the median population diameter at both pH 6.8 and pH 7.4 (**Fig 7D**), from 8.9 microns to 7.4 microns at pH 7.4 and 9.3 microns to 7.0 microns at pH 6.8. UDP-glucuronic acid, a GXM precursor, induced a significant shift in median diameter as well.

**Figure 7:**
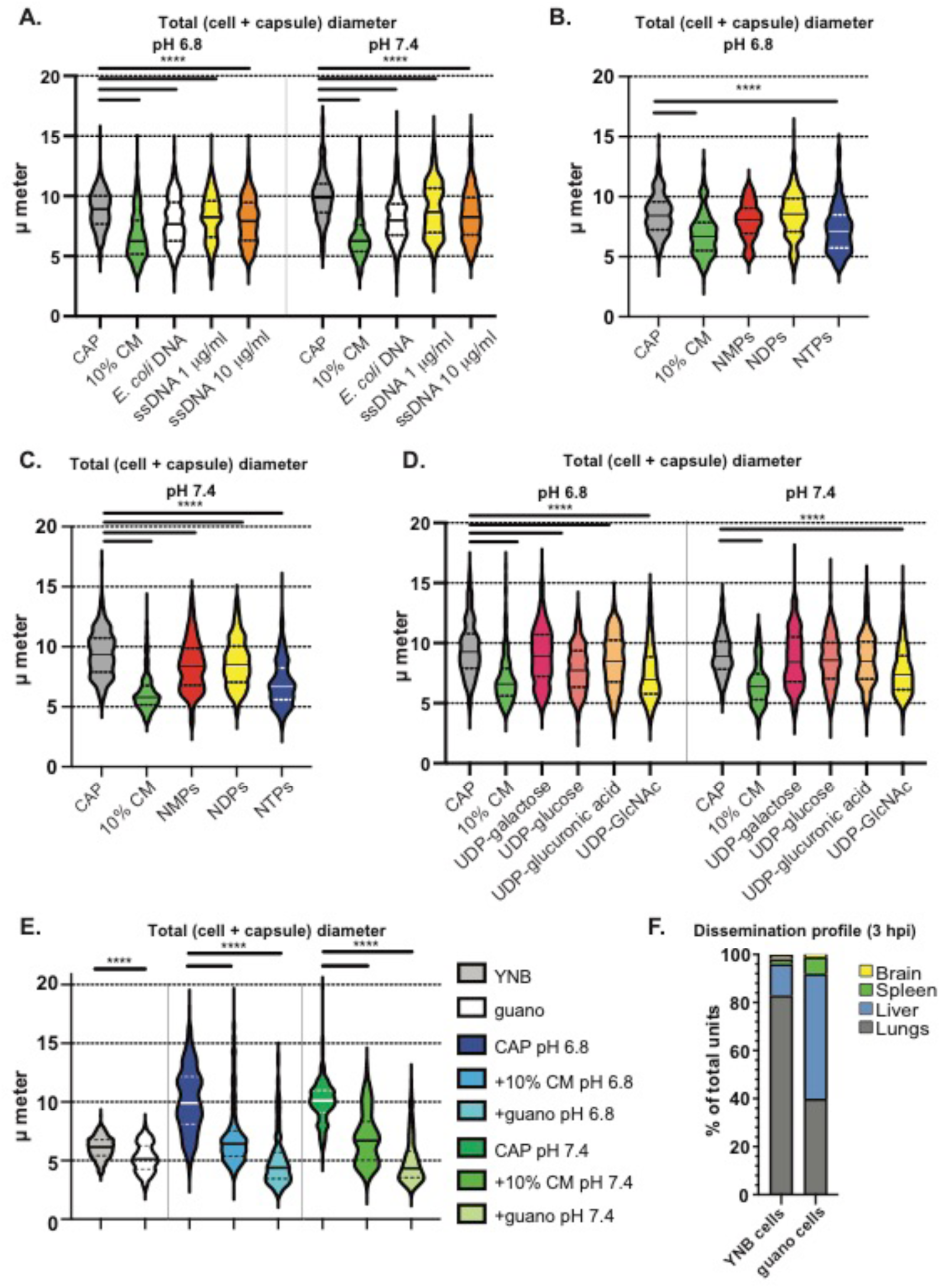
Diverse phosphate sources, including pigeon guano, induce small cell formation. **(A)** Total diameter (cell + capsule) measurements of cells after growth in CAP-medium (10% Sabouraud’s pH 6.8 (left) or 7.4 (right)) and supplementation with different DNA sources. *E. coli* DNA concentration was 1 μg/ml and ssDNA at either 1 μg/ml (yellow) or 10 μg/ml (orange). **(B)**Total diameter measurements of CAP-grown cells after exposure to nucleotide mono-, di-, or triphosphate pools at 200 μM total concentration (50 μM each of adenine, cysteine, guanine, and uridine mono-, di-, or triphosphates) following growth at pH 6.8 or (**C)** pH 7.4. **(D)** Total diameter measurements of CAP-grown cells after exposure to UDP sugars at pH 6.8 (left) or pH 7.4 (right). **(E)** Total diameter measurements of CAP-grown cells after exposure to pigeon guano extract medium. **(F)** Organ distribution following tail vein infection of *C. neoformans* cells grown in either YNB or pigeon guano extract medium. Panels **A-E**: P-values calculated using Mann-Whitney test; N = 3 biological replicates per experiment and >100 cells measured per replicate. The solid line represents the population median and the dotted lines the 25^th^ and 75^th^ percentiles. The solid line represents the population median and the dotted lines the 25^th^ and 75^th^ percentiles. All comparisons marked as significant represent populations for which the 95% confidence interval does not overlap. ****p<0.0001.

However, the largest shift in cell size we observed was in response to pigeon guano (**Fig. 7E**, **Fig. S6C,D**), which surpassed even conditioned medium as an inducing agent. The median total diameter in CAP at pH 6.8 was reduced from 9.9 microns to 4.4 microns with exposure to 10% guano medium. At pH 7.4, guano medium induced a shift in median total diameter from 10.1 microns to 4.3 microns. Since volume is a function of the radius cubed (V = 4/3 x π x r^3^) and radius = ½ x diameter, the volume of capsule-induced cells is 11-fold greater than the volume of a guano-induced small cell. Even relatively modest reductions in diameter, such as that induced by UDP-GlcNAc at pH 7.4, represents an almost 2-fold reduction in volume compared to CAP-grown cells.

In addition to inducing a substantial change in size and volume, growth in pigeon guano extract medium changes organ entry abilities of *C. neoformans* cells. We found that growing cells in 10% guano medium, even without inducing small cell formation, resulted in more entry into extra pulmonary organs than YNB-grown cells (**Fig. 7F**). Together, these data demonstrate that *C. neoformans*’s environmental niche can make *C. neoformans* cells more prone to dissemination and extrapulmonary organ entry and demonstrates the importance of considering environmental niches when studying these opportunistic pathogens.

## DISCUSSION

Here we demonstrate the formation of a new inducible morphotype that readily enters and survives in extrapulmonary organs. Since these cells are transcriptionally distinct (**Fig. 5**) and formed in response to extracellular signals (**Fig. 5, 6, 7**), we argue that they represent a separate morphotype rather than part of a continuum of variously sized cells. Given their importance for dissemination and extrapulmonary organ invasion and proliferation, we suggest the name “seed cells” for this morphotype.

While seed cells are far from the only smaller-sized morphotype in *C. neoformans*, there are notable distinct differences between seed cells and other morphotypes. Microcells, at approximately 1 micron in diameter, are smaller than seed cells (Feldmesser et al., 2001). Drop cells, while similar in shape, are described as metabolically inactive (Alanio et al., 2015a), which seed cells are decidedly not. Cell division is required for small cell formation *in vitro*, since if we do not add fresh growth medium when inducing small cells *in vitro*, the median population diameter does not shift (**Fig. 6B**). One possibility is that seed cells and Titan daughter cells (“titanides”) (Zhou et al., 2020) are the same, independently described morphotype. We cannot completely eliminate the possibility *in vivo*, but a key difference *in vitro* is that seed cell formation does not require Titan cells. We can obtain small cells *in vitro* without inducing titan cells beforehand (**Fig. 6**). In other ways, titanides are similar to seed cells, particularly their hypothesized role in dissemination. Seed cells are also about the same size as “drop” cells (Alanio et al., 2015b). These cells were isolated due to high non-cytoplasmic levels of the glutathione stain CMFDA, which measures oxidative stress response. Drop cells are not thought to be viable (Alanio et al., 2015b), while seed cells require nutrients for induction.

This work and others’ underscore the critical importance of phenotypic heterogeneity in the dissemination process. Heterogeneity in the form of antigen or phase switching is common in bacteria (van der Woude and Bäumler, 2004) and parasite infections (Deitsch et al., 1999; Ward et al., 1999). *C. neoformans* and other fungi exhibit incredible morphotype diversity but do not, to our knowledge, exhibit classic antigenic switching. The closest comparable example could be the adhesin *EPH* gene family regulation in *Candida glabrata* (López-Fuentes et al., 2018), a diverse family of subtelomeric genes whose epigenetic regulation bears some resemblance to the *var* gene family of *Plasmodium falciparum* (Kim, 2012). Instead, fungi shift their surface antigens in response to environmental changes (Ballou et al., 2016; Hommel et al., 2018; Trevijano-Contador et al., 2018) or through morphological changes (Fernandes et al., 2018) such as spore (Botts et al., 2009), titan cell (Okagaki et al., 2010), drop cells (Alanio et al., 2015a), micro cell (Feldmesser et al., 2001) and seed cell formation in *C. neoformans*.

Changes to mannose exposure on the *C. neoformans* cell surface seems to be one of the more important changes for seed cells’ interactions with the host immune system. Prior data supporting the importance of mannose recognition and mannose-binding lectin by the host during cryptococcosis is contradictory (Eisen et al., 2008; Fang et al., 2015). Despite the increased mannose exposure on small *ex vivo* cells, we did not see significant MBL-binding to cryptococcal cells. This may be because mannose-binding lectin is abundant in serum but we analyzed fungal cells from perfused lung tissue. We found increased surfactant protein D (SPD) binding of small and mid *ex vivo* cells compared with large *ex vivo* cells (**Fig. 3A**), and SPD can bind to mannose-rich microbial features (Sahly et al., 2002). Mannose receptor-deficient mice are more susceptible to *C. neoformans* infection (Dan et al., 2008). However, mannose recognition via the mannose receptor was dispensable for murine, but not primary human, macrophage phagocytosis of *C. neoformans and C. gattii* cells *in vitro* (Lim et al., 2018). Mannose recognition depends on a given fungal cell’s morphotype: the addition of exogenous mannan reduced macrophage association with seed cells, but not large *ex vivo* cells or small YNB-grown cells. Increased mannose exposure may result in a balance between host and microbe: in places such as the liver, increased mannose-dependent uptake (**Fig. 3G**) is countered by robust seed cell growth within the organ and the attempted fungal sequestration backfiring on the host (**Fig. 3F**).

Our data also indicate that growth conditions dramatically influence dissemination ability independent of size. Namely, cells originating from more stressful environments such as host lungs or host-like *in vitro* conditions were better primed for survival and dissemination *in vivo*. Capsule formation was not sufficient to facilitate organ entry (**Fig. 4A**) but did allow modest increase in blood survival (**Fig. 4E**), while YNB-grown cells were deficient in both organ entry and blood growth. These data suggest that efficient organ entry requires that cryptococcal cells adopt smaller cell and capsule morphologies (**Fig. 4A**), stress resistance that will prime them for survival in the blood and tissue (Alanio et al., 2015b; Ngamskulrungroj et al., 2012), and cell surface changes.

Nutritional immunity is an important aspect of the host-pathogen arms race. This has classically been associated with transition metals such as iron (Hood and Skaar, 2012). However, phosphate availability is also tied to virulence (Kretschmer et al., 2014; Lev et al., 2017). Here we find that phosphate is sufficient to induce seed cells at pHs where phosphate is more likely to be limiting (**Fig. 6**).

Phosphate is essential for cell homeostasis and growth, and its acquisition is tightly controlled in microbes that absorb fluctuating levels of phosphate from their surroundings (Kohler et al., 2020; Lev and Djordjevic, 2018). Phosphate is critical for *Candida albicans* morphogenesis (hyphal formation), stress resistance, and tissue invasion (Ikeh et al., 2016; Liu et al., 2018). Phosphate acquisition is also essential for survival in the blood and dissemination to the brain (Lev et al., 2017). *Cryptococcus* relies on a proton gradient to import phosphate (Lev and Djordjevic, 2018) and therefore survives poorly in the alkaline (pH 7.4) bloodstream where its ability to upregulate phosphate acquisition machinery is impaired (Lev et al., 2017). Our results suggest that increased phosphate availability also impacts dissemination at an even earlier stage by mediating the emergence of seed cells within the lungs (**Fig. 7**). We observed an order of magnitude decrease in lung pH over the course of infection, which could also enhance acquisition of phosphate and other nutrients (Lev et al., 2017) or independently repress capsule growth (Farhi et al., 1970; O’Meara and Alspaugh, 2012) while still permitting seed cell formation.

*PHO4* is not necessary for seed cell formation at pH 6.8. There could be at least two pathways that control seed cell formation, with one phosphate-induced and one occurring at acidic pH. Alternatively, the rewiring of the Pho4-controled genetic network at acidic pH removes Pho4-dependence of seed cell formation-driving genetic factors. These formation signals might or might not be present at both acidic and alkaline pH – pigeon guano and conditioned medium both still induce seed cell formation at a range of pHs – but this will be a fruitful area for future investigation.

Another source of variation within a population, which is sometimes linked to morphology, is fungal cell ploidy. Ploidy serves as a driver of adaptation, in-host survival, and pathogenicity in multiple fungal species (Gerstein et al., 2015; Okagaki and Nielsen, 2012; Selmecki et al., 2015). Of our *ex vivo* populations, only large cells contain >2N cells (**Fig. S2**), and even within the large cell population, the majority of cells are 2N, which could be either cells in the G2 phase of mitosis or smaller titan cells. Intermediate cells display the DNA content profile of actively growing haploid yeast cells (Todd et al., 2018). We therefore think that comparisons between seed and intermediate *ex vivo* cells are likely independent of ploidy.

There are some notable limitations to our study. For example, we did not initially define the subpopulations within the lungs based on clear biological criteria, but rather sorted cross-sections of the population of fungal cells proliferating within the lungs. Small *ex vivo* “seed” cells correspond with the size of disseminated cells in the brain, but intermediate and large cells are based on arbitrary size cutoffs designed to distinguish them from small cells. Clearer definitions of fungal subpopulations within the lungs and analysis of their dissemination would enhance our understanding of systemic cryptococcosis. Furthermore, *C. neoformans* faces multiple bottlenecks during dissemination, including escaping the lungs. *C. neoformans* can escape the lung mucosa to access the bloodstream by crossing lung epithelial cells or via the Trojan horse mechanism within macrophages (Denham and Brown, 2018). We focused on extrapulmonary organ entry rather than examining lung escape by seed cells directly. However, seed cells are more likely to be phagocytosed, which may lead to increased Trojan horse dissemination. We also find that in late infection (17 dpi), the total diameter of cells in the blood shows a size distribution similar to the lungs (**Fig. 1E**), suggesting that pulmonary escape may be less of a bottleneck than previously thought.

Phenotypic heterogeneity is an important microbial property that facilitates severe, disseminated infections (Zaragoza, 2011; Zhou and Ballou, 2018). Here we demonstrate that an individual morphotype, the seed cell, is prone to disseminate and shows an increased ability to enter extrapulmonary organs. Seed cells’ dissemination ability is consistent with observations in humans (Fernandes et al., 2018), in that phenotypic heterogeneity strongly influences clinical outcome. Small *C. neoformans* cells are also found in the brains of human patients with cryptococcal meningitis (Xie et al., 2012). Overall, seed cells represent a new inducible morphotype formed in response to phosphate and other signals. Infection-induced phosphate availability could establish a feed-forward loop that results in self-propagation of *C. neoformans* dissemination. Moreover, conditions in *C. neoformans*’s environmental niche can induce seed cell formation and increase organ entry, thus increasing disease-causing potential for even an “accidental” pathogen.

## ACKNOWLEDGEMENTS

This work was supported by research grant R01AI130248 from the National Institutes of Health to J.C.S.B. S.T.D. was supported by grant T32AI055434 from the National Institutes of Health. Flow cytometry work was supported by the University of Utah Flow Cytometry Facility through the National Cancer Institute Award Number 5P30CA042014-24 and by the National Center for Research Resources of the National Institutes of Health under Award Number 1S10RR026802-01. RNA sequencing and analysis was performed by the High-Throughput Genomics and Bioinformatics Analysis Facility, a part of the Huntsman Cancer Institute at the University of Utah. Generation of the *pho4*Δ #1 clone was constructed in the Madhani laboratory with support from National Institutes of Health funding (R01AI100272). We would also like to thank Dr. Kyla Ost for her valuable input regarding the analysis of opsonin-binding via flow cytometry. We would also like to thank Dr. Matthew Rondina and Dr. Alicia Eustes for their assistance with *in vivo* platelet depletion.

## AUTHOR CONTRIBUTIONS

S.T.D. and J.C.S.B. conceived and designed the experiments. S.T.D., B.B., K.Y.C., M.A.W., J.M.B., and J.C.S.B. performed the experiments. S.T.D., B.B., and J.C.S.B. analyzed the data. S.T.D. and J.C.S.B. wrote the paper. S.T.D., B.B., K.Y.C., M.A.W., and J.C.S.B. edited the paper.

## METHODS

### Fungal strains and growth media

The following strains were used in this study: KN99α (Brown lab stock which was also the background for mutant strains), Bt63 (Brown lab stock, NCBI:txid1295841), 52D (ATCC 24067), R265 (ATCC MYA4093), WM276 (ATCC MYA-4071).

The following growth media were used in this study: YNB medium (yeast nitrogen base without amino acids [Difco catalog no. 291940], 2% glucose), CAP medium (10% Sabouraud’s dextrose [Difco catalog no. 238230], buffered with 50 mM HEPES (pH 7.4) or MES (pH 6.3)), Titan cell medium (5% Sabouraud’s dextrose, 10% fetal calf serum (GenClone catalog no. 25- 550), 15 μM sodium azide, 50 mM HEPES (pH 7.4)).

### *In vitro* small cell induction assay

To screen for factors that facilitated shifts in cell size, we picked single colonies from *C. neoformans* cells streaked on YPAD agar and cultured them overnight (12-18 hours) in YNB medium at 30 °C. We then sub-cultured those cells into YNB medium, CAP medium, or titan cell medium for 24 hours at an initial cell density of 10^5^ cells/mL. YNB- and CAP-grown cells were cultured at 37 °C, while titan cell medium-grown cells were cultured at 37 °C and 5% CO2. At this stage, conditioned medium (CM) was collected from the 24-hour YNB cultures by pelleting the fungal cells and filtering (0.22 μm pore size) the supernatant. 450 μl of the CAP and titan cell media cultures were sub-cultured into 450 μl fresh media and 50 μl (10% of the final volume) of one of the following supplements solubilized in H2O: 10% H2O (vehicle control), CM, 1% CM, 50 μg GXM, 50 μM Qsp1 peptide (Peptide 2.0: NFGAPGGAYPW), 50 μM Qsp1 scrambled peptide (Peptide 2.0: AWAGYFPGPNG), 10% 1x PBS (phosphate-buffered saline), 10% YNB medium; the following YNB medium supplements were added to a final concentration that equaled their concentrations in YNB medium: 2% glucose, 4.531 nM folic acid, 8.186 nM biotin, 250.6 nM copper sulfate pentahydrate, 531.4 nM riboflavin, 602.4 nM potassium iodide, 839.5 nM calcium pantothenate, 971.3 nM Sodium molybdate dihydrate, 1.186 μM thiamine hydrochloride, 1.233 μM ferric chloride hexahydrate, 1.458 μM p-aminobenzoic acid, 1.945 μM pyridoxine hydrochloride, 2.477 μM zinc sulfate heptahydrate, 2.649 μM manganese sulfate monohydrate, 3.249 μM niacin, 8.087 μM boric acid, 11.10 μM inositol, 901.1 μM calcium chloride dihydrate, 1.711 mM sodium chloride, 4.154 mM magnesium sulfate heptahydrate, 7.348 mM potassium phosphate monobasic, and 37.8mM ammonium sulfate. The supplemented cultures were incubated for another 24 hours at 37 °C, following which 400 μl was aliquoted to estimate cell size by flow cytometry and another 400 μl was aliquoted to measure cell and capsule size (see “fungal cell size measurements”) in India ink (Higgins catalog no. 44201).

### Pigeon guano medium

Pigeon guano was a gift from Michael Shapiro’s lab (University of Utah Department of Biology). Following collection, it was lyophilized, then stored at room temperature until use. Lyophilized guano was ground in a coffee grinder until a powder. Guano medium consisted of a 20% w/v solution in water, which was boiled for 10 minutes, then filtered through first Whatman 3 mm filter paper, then a 0.5 μm polystyrene filter, and finally a 0.2 μm polystyrene filter. Resultant medium was stored at room temperature for up to four weeks.

### Phosphate source small cell induction assay

We grew fresh colonies of KN99 cells overnight in YNB at 37°C, then subcultured at 10^5^ cells/ml in CAP-medium 10% Sabouraud’s buffered to either pH 7.4 with 50 mM HEPES or pH 6.8 with 50 mM Tris HCl). Cultures were grown 24 hours at 37°C, then diluted 1:1 with either 20% (w/v) pigeon guano medium or fresh 10% Sabouraud’s of the appropriate pH containing the phosphate source of interest (DNA; nucleotide mono-, di-, or triphosphates; or UDP sugars).

### Mouse infection models and fungal burden analysis

For intranasal inoculations, ∼8-week-old female C57BL/6NJ mice (Jackson Laboratory) mice were anesthetized with ketamine/dexmedetomidine hydrochloride (Dexdomitor) delivered intraperitoneally. They were then suspended by their front incisors on a horizontal strand of thread. Unless otherwise indicated, mice were inoculated intranasally with 2.5x10^4^ *Cryptococcus* cells in 50 μl of 1XPBS using a micropipette. The inoculum was placed dropwise onto a nasal flare before being inhaled by mice. Ten minutes later, mice were intraperitoneally administered the reversal agent atipamezole (Antisedan).

For intravenous inoculations, ∼8-week-old female C57BL/6NJ mice (Jackson Laboratory) were warmed under a heat lamp before being placed in a restraint. Mice were inoculated via the tail vein with 10^5^ *C. neoformans* cells or beads in 200 μl of 1XPBS using 28- gauge x 12.7 mm syringes. In order to competitively inhibit host interactions with fungal components *in vivo*, we administered 400 µg of GXM (see “GXM isolation), mannan (Sigma-Aldrich catalog no. M7504), β-glucan (Millipore Sigma catalog no. 1048288), the chitin monomer N-acetyl glucosamine (Vector Laboratories S-9002), or polyinosinic acid (Poly(I)) (Sigma-Aldrich catalog no. 26936-41-4) in 200 μl of 1XPBS 2-3 minutes prior to inoculation with fungal cells.

For intracranial inoculations, ∼6-week-old female C57BL/6NJ mice (Jackson Laboratories) were anesthetized with ketamine/dexmedetomidine hydrochloride as described above. They were inoculated intracranially with 10^3^ *C. neoformans* cells in 30 μl of 1XPBS via a 26-gauge 1/2-inch needle. Following inoculation, mice were intraperitoneally administered the reversal agent atipamezole (Antisedan; ∼0.0125 mg/g).

Nonperfused mice were euthanized by CO2 asphyxiation and cervical dislocation. Mice that were intracardially perfused at time of death were anesthetized with isoflurane and perfused in a nonrecirculating fashion before cervical dislocation. Fungal burden was assessed by excising organs and homogenizing them in 5 mL 1XPBS, washing the probe in between samples: 30 seconds 10% bleach, 45 seconds 70% EtOH, and 10 seconds sterile H2O. Ten-fold serial dilutions of organ homogenate were plated on Sabouraud’s agar containing 10 mg/ml gentamicin and 100 mg/ml carbenicillin and stored at 30°C for 3 days before counting CFUs.

All animal procedures were approved by the University of Utah Institutional Animal Care and Use Committee.

### Fungal cell size measurements

*C. neoformans* cells harvested from laboratory growth medium or infected mouse lungs were fixed in 2% paraformaldehyde for 20 minutes. To visualize capsule, 4 μl of India ink with 4 μl of cell suspension on a microscope slide. At least 10 successive images were taken starting at one edge of the coverslip and moving across to the opposite side, as smaller cells tended to drift to the edge of the coverslips. Total diameter was measured from one edge of the capsule to the other. Cell body diameter was measured from one edge of the cell wall to the other. Capsule thickness was calculated as follows: capsule thickness = (total diameter – cell body diameter) /2 The total number of cells counted for a given experiment is indicated in the figure legends.

### FACS isolation of fungal cells

Mice were intranasally inoculated with *C. neoformans* KN99-mCherry (see “Mouse infection models”). At 14 dpi, lungs from 3 mice were excised and placed in 15 mL of 1XPBS. The three lungs were homogenized together using a mechanical tissue homogenizer. The homogenate was filtered first through a 70 μM cell strainer and then through a 40 μM cell strainer to prevent downstream clogging of the flow cytometer. Cells were centrifuged at 2195xg for 10 min. The supernatant was discarded and the pellet was resuspended in ∼1mL 1XPBS. The suspension was filtered once more through a 70 μM cell strainer and was resuspended at ∼1x10^7^ cells/mL in 1XPBS for FACS (BD FACS Aria). The FACS gating scheme is represented in **Figure 2**.

### Isolation of polystyrene beads from mouse organs

Mice were intravenously administered (see “Mouse infection models”) 10^5^ green fluorescent polystyrene beads (Polysciences, Inc. catalog no. 17156-2 and 18241-2), and 3 hours later the indicated organs were harvested. Organs were homogenized with a mechanical tissue homogenizer in 5 mL tissue lysis buffer (100 mM Tris-Cl pH 8.0, 200mM NaCl, 5mM EDTA, 0.2% (w/v) SDS), washing the probe in between samples: 15 seconds H2O, 15 seconds 70% EtOH, and 15 seconds fresh H2O.

Beads from each organ were concentrated via differential centrifugation. The homogenates were poured into a fresh 50 mL conical. The homogenization tubes were washed with 30 mL of H2O and combined with the respective homogenized sample. The samples were filtered through cell strainers (70 μM pore size), after which another 10 mL of H2O was passed through each strainer. The samples were centrifuged at 4480xg for 40 minutes. The supernatant was aspirated to the 5 mL mark and the samples were transferred to fresh 15 mL tubes containing 2 mL of 100% Percoll (GE Healthcare catalog no. 17-0891-01). Water was added to the 10 mL mark. The tubes were vortexed to mix and allowed to settle for 5 minutes before centrifugation at 1120xg for 45 minutes. The supernatant was aspirated to the 2 mL mark. 10 mL of H2O was added to the samples before centrifuging at 4480xg for 20 minutes. The supernatant was aspirated and resuspended in 200-1000 μl 1XPBS for flow cytometry. Entire samples were analyzed to determine the number of beads per organ.

### Fungal cell staining and flow cytometry analysis

Between 10^5^ and 10^6^ *C. neoformans* cells harvested from laboratory growth medium or infected mouse lungs were pelleted and fixed in 2% paraformaldehyde for 20 minutes. Fixed cells were pelleted and washed twice with 1XPBS. Cells were then resuspended in 1XPBS and stained with the desired antibodies/lectins/fluorescent dye.

To estimate DNA content, cells were stained for 10 minutes (room temperature) in 0.3 μg/mL 1XPBS+0.1% triton X-100. Mid-log phase fungal cells grown in YNB medium +/- 80 μg/mL benomyl (Agilent catalog no. PST-1245) for 24 hours were used to set 1C and 2C gates. Untreated mid-log phage fungal cells display two DAPI peaks (1C ad 2C), while benomyl prevents cell division and traps cells at 2C.

To estimate fungal microbial associated molecular pattern exposure, cells were stained at room temperature with 5 μg/mL fluorescein-conjugated wheat-germ agglutinin (WGA, Vector Laboratories catalog no. FL-1021) for 15 minutes to detect exposed chitin or 50 μg/mL fluorescein-conjugated concanavalin A (ConA, Vector Laboratories catalog no. FL-1001) for 5 minutes to detect exposed mannose.

To estimate the percentage of cells bound by soluble host factors, we used the following antibodies: α-complement C3 (primary: mAb 11H9, ThermoFisher catalog no. MA1-40046; secondary: Mouse anti-Rat IgG2a, FITC, Thermofisher catalog no. 11-4817-82), α-surfactant protein D (primary: polyclonal, Abcam catalog no. ab203309; secondary: Donkey anti-Rabbit IgG, AlexaFluor 405, Abcam catalog no. ab175651), α-mannose binding lectin A (primary: mAb 8G6, Hycult catalog no. HM1035; secondary: Mouse anti-Rat IgG2a, FITC, Thermofisher catalog no. 11-4817-82), α-mannose binding lectin C (primary: mAb 14D12, Abcam catalog no. ab106046; secondary: Mouse anti-Rat IgG2a, FITC, Thermofisher catalog no. 11-4817-82), α- IgM (mAb II/41, FITC, BD Biosciences catalog no. 553437), α-IgA (mAb 11-44-2, FITC, SouthernBiotech catalog no. 1165-02). Cells were stained with primary antibodies (1:50 dilution) for 30 minutes on ice. When a secondary antibody was required, primary antibody labeled cells were pelleted, washed twice with 1XPBS and stained with the secondary antibody (1:50 dilution) for 20 minutes on ice.

After staining, cells were pelleted and washed twice with 1mL 1XPBS before being resuspended in 200-400 μl of 1XPBS for flow cytometry.

### GXM isolation

GXM was isolated as described previously (Wozniak and Levitz, 2009) with slight modifications. *C. neoformans* (KN99) cells were grown in 100 mL for 3 days at 37°C in YNB medium (YNB+2% glucose). Cells were centrifuged for 15 minutes at 12,000xg. The supernatant was filtered (0.45 μM pore size) and 50 mL of supernatant was transferred to a 250 mL bottle. ∼150 mL of 95% EtOH was added to the samples in 4-5 aliquots, mixing after each addition. Samples were stored at 4°C overnight to precipitate GXM. Samples were centrifuged for 30 minutes at 15,000xg, 4°C. The pellets were resuspended in 0.2M NaCl to a concentration of about 10 mg/mL (∼50 mL per 250 mL bottle). Samples were transferred to a clean beaker stirred until samples were completely resuspended. 3 mg hexadecyltrimethylammonium bromide (CTAB) per 1mg precipitate was slowly added with stirring and low heat. Once solubilized, the samples were removed from heat and 250 mL of 0.05% CTAB was slowly added with stirring. The samples were moved to fresh 250 mL bottles and centrifuged at for 2 hours at 11,000xg, 4°C. The supernatant was discarded and the pellet was washed in 150 mL of 10% EtOH. The samples were centrifuge for 20 minutes at 15,000xg, room temperature. The supernatant was discarded and the pellets were resuspended in 50 mL of 1M NaCl. The samples were transferred to a clean beaker and stirred until completely resuspended.

Approximately 2 volumes of 95% EtOH was added dropwise with stirring to precipitate the GXM while leaving the CTAB in solution. The samples were centrifuged in fresh 250 mL bottles for 20 minutes at 11,000xg, room temperature. The supernatant was discarded and the precipitate was dissolved in 2M NaCl. The resulting solution was placed in a snake-skin dialysis cassette with a 3,500 molecular weight cutoff (Thermo Fisher Scientific catalog no. 68035) and dialyzed overnight against 1M NaCl. The samples were subsequently dialyzed against distilled H2O for 14 days at 4°C. The distilled H2O was changed every 1-2 hours for the first 2 days, and then 4-5 times a day for the next 6 days. For the last 2-6 days, the water was changed 2-3 times a day. The dialyzed samples were lyophilized and stored at -80 °C.

### Cell-type specific depletion in mice

Hepatic and splenic macrophages were depleted from mice using clodronate liposomes. Mice were intravenously administered 200 µl of clodronate liposomes (Liposoma BV catalog no. C-010) or PBS control liposomes (Liposoma BV catalog no. P-010) 48 hours prior to inoculation with *C. neoformans* cells. Macrophage depletion efficiency in the liver and spleen was assessed on the day of inoculation via flow cytometry. Macrophages were identified as CD45+ (anti-CD45-efluor450, eBiosciences catalog no. 48-0451-82) and F4/80+ (anti-F4/80-APC, eBiosciences catalog no. 17-4801-80).

Neutrophils were depleted from mice using anti-Ly6G antibodies. Mice were intraperitoneally administered 200 ug of anti-Ly6G (clone 1A8 BioXcell catalog no. BP0075-1) antibodies or PBS control at 24 and 2 hours prior to inoculation with *C. neoformans* cells. Neutrophil depletion efficiency in the blood was assessed on the day of inoculation via flow cytometry. Neutrophils were identified as CD45+ (anti-CD45-efluor450, eBiosciences catalog no. 48-0451-82), CD11b+ (anti-CD11b-APC, eBiosciences catalog no. 17-0112-82), and Ly6G+ (anti-Ly6G-FITC, eBiosciences catalog no. 11-5931-82).

Platelets were depleted from mice using anti-GPIbα antibodies. Mice were intravenously administered 80 μl of anti-GPIbα antibodies (Emfret catalog no. R300) or PBS control 24 hours prior to inoculation with *C. neoformans* cells. Platelet depletion efficiency was determined by Hemavet 950FS (Drew Scientific Group) analysis of blood collected by cheek bleed 10 minutes prior to anti-GPIbα administration and on the day of inoculation.

### Liver macrophage isolation and infection

Liver macrophages were isolated from 8- to 10-week-old female C57BL/6NJ mice (Jackson Laboratory) according to established procedures (Li et al., 2014). Isolated liver macrophages were resuspended in complete DMEM (Dulbecco’s Modified Eagle’s Medium, GenClone catalog no. 25-500) supplemented with 10% fetal bovine serum (FBS, (GenClone catalog no. 25-550) and 100 U/mL Penicillin/Streptomycin and seeded into a T-25 flask at a density of 8-10 × 10^6^ cells/flask. Macrophages were allowed to settle and adhere for 4 hours in a mammalian tissue culture incubator (37 °C, 5% CO2). Non-adherent cells were then removed from the dish by gently washing 3 times with 1XPBS (without divalent cations), leaving adherent macrophages. The media was replaced with 5 mL complete DMEM, and the cells were rested for 4 days. The media was replaced on the day after initial seeding, and then as needed.

Fungal infection and association with liver macrophages was assessed using the following procedures. The media was aspirated from rested liver macrophages and the cells were washed once with 1XPBS. Macrophages were lifted from the plate by treating them with 1mL Accutase (Corning catalog no. 25-058-CI) for ∼5 minutes, tapping the plate to dislodge cells. 1 mL complete DMEM was added to the Accutase-treated cells, which were then centrifuged at 300xg for 5 minutes. Liver macrophages were resuspended cells in 0.5-1 mL complete DMEM and live cells were counted by mixing trypan blue (HyClone catalog no. SV30084.01) with cell suspension at a 1:1 ratio and counting clear cells on a hemocytometer. 40,000 live cells/well were seeded into 96-well plates in complete DMEM. The liver macrophages were allowed to settle and adhere overnight.

The next day, the inoculum was prepared by resuspending *C. neoformans* cells in 500 μl of 0.01% Direct Yellow 96 (AK Scientific 61725-08-4), vortexing briefly to mix, and incubating for 5 minutes on the benchtop. *C. neoformans* cells stained with Direct Yellow 96 were pelleted, washed once with 1 mL 1XPBS, and resuspended at a concentration of 8x10^6^ cells/mL in serum-free DMEM. The supernatant was aspirated from the macrophages seeded in the 96-well plates and was replaced with 200 μl of serum-free DMEM. 10 μl of the inoculum (8x10^4^ cells) was added to each well, resulting in a multiplicity of infection of 1:1. Macrophages and *C. neoformans* cells were co-cultured for 4 hours in a mammalian tissue culture incubator (37 °C, 5% CO2). The medium was then aspirated from each well, and the cells were washed three times with 100 μl of 1XPBS to remove non-macrophage associated fungal cells. The remaining cells were fixed in 100 μl of 4% paraformaldehyde for 10 min. The paraformaldehyde was then removed and the cells were washed twice with 100 μl of 1XPBS and stained with100 μl of DAPI (Sigma Aldrich catalog no. 28718-90-3) working solution (600 nM DAPI; 0.1% Triton X-100; in 1XPBS) for 5 minutes. The DAPI working solution was then removed and cells were washed twice with 100 μl of 1XPBS. Finally, 100 μl of 1XPBS was added to the wells to prevent desiccation. Cells were imaged on a Nikon widefield microscope, and the number of macrophages with associated *C. neoformans* cell(s) were counted (green-fluorescent due to Direct Yellow 96). Six images were taken per well in the same relative positions and every imaged macrophage was scored as having a physically associated fungal cell(s) or not, resulting in 2263-3197 macrophages counted per experimental condition across three independent replicates.

In order to asses liver macrophage killing of *C. neoformans* cells, liver macrophages were seeded in 96-well plates and infected with *C. neoformans* cells as described above. The plates containing *C. neoformans* and macrophages were incubated for 24 or 48 hours. After the desired time, the supernatant was collected from each well and transferred to a microcentrifuge tube. 200 μl of sterile distilled water was then added to each well in order to lyse the macrophages for 40 minutes at 37°C. The contents of each well were then mixed to collect every *C. neoformans* cell and combined with the respective supernatant now in a microcentrifuge tube. The wells were rinsed with 200 μl sterile PBS and added to the respective microcentrifuge tube. Serial dilutions of each of the well contents were plated on YPAD agar and incubated at 30°C for 2-3 days before counting CFUs. CFUs quantified at each timepoint were normalized to the CFUs in the inoculum in order to calculate the “% of input”. Each macrophage killing experiment was repeated on three different days with each experimental condition being performed in duplicate wells.

### Fungal survival in blood

We collected whole blood from 8- to 10-week-old female C57BL/6NJ mice (Jackson Laboratory) to assess *C. neoformans* survival in blood. Mice were anesthetized with isoflurane, and whole blood was collected by cardiac puncture using 28-gauge x 12.7 mm syringes pre-coated with 0.5M EDTA. Blood was immediately transferred to tubes containing heparin to achieve a final concentration of 30 units of heparin per mL of blood. 100 μl of blood was added to individual wells of a 96-well plate. 100 *C. neoformans* cells in 5 ul of 1XPBS were used to inoculate each well. The plates were incubated in a mammalian tissue culture incubator (37 °C, 5% CO2) without shaking. At the indicated timepoint, the well contents were transferred to fresh microcentrifuge tubes. The wells were washed with 100 ul of 0.05M EDTA, which was then added to the respective microcentrifuge tube. The entire contents of the microcentrifuge tubes were plated on YPAD agar and incubated at 30°C for 2-3 days before counting CFUs. CFUs quantified at each timepoint were normalized to the CFUs in the inoculum in order to calculate the “% of input”. Each experiment was repeated on three different days with each experimental condition being performed in duplicate wells.

### RNA isolation, sequencing, and analysis

RNA sequencing data is publicly available at NCBI GEO (Accession number: GSE152784). *C. neoformans* cells were harvested from laboratory media and from infected mouse lungs for RNA isolation and sequencing. Three replicates were sequenced for each sample with replicates being defined as follows. Approximately 10^7^ cells were harvested from the indicated *in vitro* growth condition on three separate days. At least 10^6^ *ex vivo* cells were isolated per sort gate each day (see “FACS isolation of fungal cells”). *Ex vivo* cells were isolated from the pooled lung homogenate of three mice. *Ex vivo* sample populations were isolated on two consecutive days and pooled for RNA isolation and sequencing, with the end result being that each sequenced *ex vivo* replicate represents fungal cells isolated from pools of six mice.

RNA was isolated using phenol-chloroform extraction, followed by Qiagen RNeasy column-based cleanup. Fungal cells were pelleted, flash frozen in liquid nitrogen, and lyophilized until completely dry. Lyophilized cells were resuspended in 750 μl of TRIzol (ThermoFisher Scientific catalog no. 15596026) inside screw-cap microcentrifuge tubes. Approximately 50 µl of 1-mm and 150 µl of 0.5mm zirconium beads (BioSpec Products catalog no. 11079110z and 11079105z) were added to each sample and placed in a Biospec Products mini bead beater to lyse (12 x 2 min beating pulses, using sample blocks stored at -20 °C to prevent overheating). 150 μl of chloroform was added to the lysed samples and incubated for 2- 3min at room temperature. Samples were centrifuged for 15 minutes at 12,000 x *g*, 4°C. The aqueous phase was transferred to a fresh tube and combined with 5-10 μg RNase-free glycogen as a carrier. 0.375 mL of isopropanol was added to each sample before incubating at least ∼20 minutes on ice. The samples were centrifuged for 15 minutes at 12,000 x *g*, 4°C. The supernatant was discarded, and the isopropanol was removed by pipetting. The samples were then briefly air-dried before resuspension in 350 μl of RLT buffer (Qiagen RNeasy) and 350 μl 70% EtOH. At this junction, samples were run through Qiagen’s RNAeasy kit (Qiagen catalog no. 74104) with on-column DNase treatment.

cDNA libraries were generated with the NEBNext Ultra II Directional RNA Library Prep with rRNA Depletion Kit (New England BioLabs catalog no. E7760) by the High Throughput Genomics Core at the University of Utah. Sample libraries were sequenced on the Illumina NovaSeq 6000. One set of samples (primarily the *ex vivo* small, medium, and large cells) were stored by the High Throughput Genomics Core at -80°C for six months prior to library construction and showed more degradation products than replicates stored at -80°C for one month. We therefore limited our analysis and interpretation of the data from these samples.

The *Cryptococcus neoformans* var. grubii H99 genome and gene feature files were downloaded from Ensembl Fungi release 46 and the reference database was created using STAR version 2.7.2c with splice junctions optimized for 150 base pair reads (Dobin et al., 2013). Optical duplicates were removed from the paired end FASTQ files using clumpify v38.34 and reads were trimmed of adapters using cutadapt 1.16 (Martin, 2011). The trimmed reads were aligned to the reference database using STAR in two pass mode to output a BAM file sorted by coordinates. Mapped reads were assigned to annotated genes in the gene feature file using featureCounts version 1.6.3 (Liao et al., 2014). The output files from cutadapt, FastQC, Picard CollectRnaSeqMetrics, STAR and featureCounts were summarized using MultiQC to check for any sample outliers (Ewels et al., 2016). Differentially expressed genes were identified using a 5% false discovery rate with DESeq2 version 1.24.0 (Love et al., 2014). Differentially expressed genes were run through FungiDB Gene Ontology (GO) Enrichment analysis, selecting for “biological process” with a p-value cutoff of 0.05 (Basenko et al., 2018; Stajich et al., 2012). Venn Diagrams were made with software from the Van de Peer group at Ghent University (http://bioinformatics.psb.ugent.be/webtools/Venn/).

### Lung pH measurements

To measure lung tissue pH, mouse lungs were excised and immediately pierced with a micro pH electrode (Orion (ThermoFisher Scientific) catalog no. 9863BN). The electrode was completely embedded in the lung tissue before taking pH readings. Two readings were taken per mouse. One reading was taken from the left lobe of the lungs and a second reading was taken from one of the right lobes. These two readings were averaged to estimate lung pH for a given mouse.

### Construction of the KN99:*pho4*Δ strains

The KN99:*pho4*Δ (*CNAG_06751*Δ) strains were constructed via homologous recombination-based replacement of the *pho4* genomic sequence with a nourseothicin resistance marker. The KN99:*pho4*Δ #1 strain was obtained from the KN99 gene deletion library (Fungal Genetics Stock Center, 2015 Madhani plates). The KN99:*pho4*Δ #2 strain was generated by PCR amplification of the nourseothicin resistance marker and ∼1,000 base pair flanking regions from the KN99:*pho4*Δ #1 strain (primers used to amplify the knock-out construct: Forward: 5’ AAAACGGCTGAAGGCTCGTTCT3’, Reverse: CTTCTGCAAGGTGAAGTTCACG). The resulting pho4 knock-out cassette was used to transform wild-type KN99 cells via biolistic transformation as described previously (Davidson et al., 2000). Putative transformants were PCR-verified by confirming the absence of the *pho4* open reading frame (Primers to verify the absence of the open reading frame: Forward: 5’CCATCTCAGATACCAACTCGCC’, Reverse: 5’CAATTTGCTGAGAGCCATAGGC3’) and the integration of the knock-out construct at the correct site (Primers to verify the correct 5’ integration site: Forward: 5’AGAGCTATATGGTATGACGAAC3’, Reverse: 5’TGTGCTGATCATCCGATGCCAC3’ and the correct 3’ integration site: Forward 5’TGTGGAGGATGGTGGGGAATAG3’, Reverse: 5’GGACGTGAGCCAATAAGTTCCT3’.

### Statistics

All statistical analyses, with the exception of RNA sequencing analysis, were performed with GraphPad Prism 8 software. Hypothesis-driven experimental comparisons were analyzed by unpaired t-test or one-way ANOVA and uncorrected Fisher’s LSD if the data held a Gaussian distribution and a Mann-Whitney U-test if the data were not Gaussian. For experiments with large sample sizes (e.g. cell size measurements), statistical significance is shown only if the 95% confidence intervals of the median compared samples does not overlap. *In vitro* experiments were replicated three times unless otherwise stated. *In vivo* (murine infection) experiments were replicated twice unless otherwise stated. Sample sizes are stated in the figure legend.

### Phosphate extraction and measure

To determine the amount of phosphate within cells, we grew cells in CAP medium and performed a standard small cell induction assay. We harvested 3 x 10^7^ total cells by centrifugation, washed briefly in water to remove any excess growth medium, then froze in liquid nitrogen. Samples were then lyophilized overnight, then resuspended in 300 μl MilliQ water, vortexed for 30 seconds on/30 seconds off (on ice) for 30 minutes, then tested for phosphate concentration using the Phosphate Colorimetric Kit (Sigma-Aldrich, catalog no. MAK030-1KT) according to the manufacturer’s instructions.

---

**Supplemental Figure 1:**
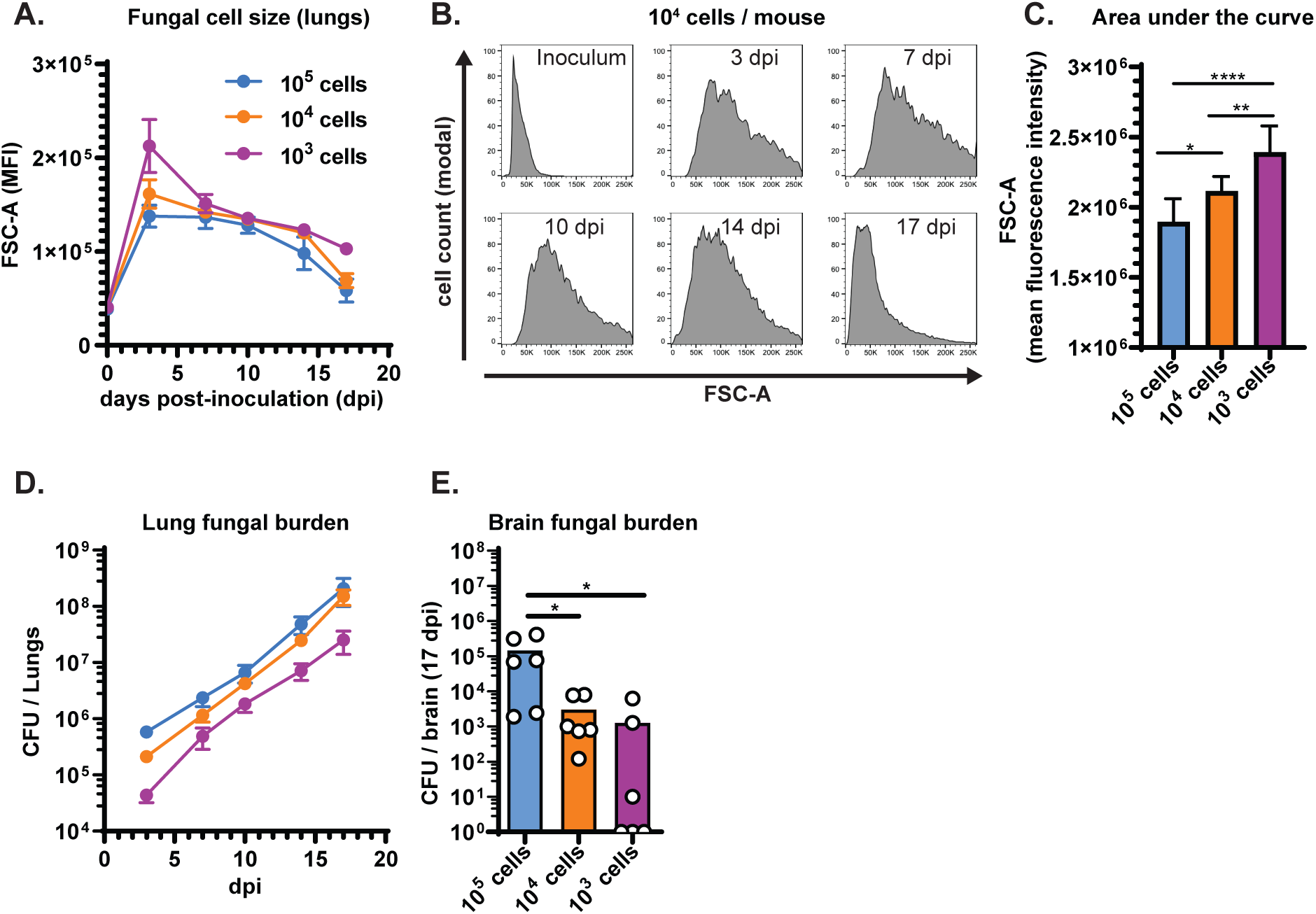
Inoculum size alters the rate at which *C. neoformans* populations shift toward smaller cell size in the lungs and disseminate to the brain. **(A)** We inoculated mice with the indicated number of KN99-mCherry cells. The line graph shows the forward scatter-area (FSC-A) measurements at the indicated days post-inoculation (dpi), which estimates relative fungal size in the lungs (N=6 mice per timepoint). Points indicate the mean and the error bars indicate standard deviation. **(B)** Representative flow cytometry plots from mice inoculated with 104 KN99-mCherry cells / mouse. **(C)** Quantification of the area under the curve for the curves displayed in Figure S1A. Bar graphs display the mean fluorescence intensity and error bars indicate standard deviation. **(D)** Lung fungal burden in the same mice described in Figure S1A and quantified by colony forming units (CFU). **(E)** Brain fungal burden in mice at 17 dpi quantified by CFUs. Bar graphs display the mean. All comparisons in Figure S1 were analyzed using one-way ANOVA and uncorrected Fisher’s LSD (ns: not significant; *p<0.05; **p<0.01; ***p<0.001; ****p<0.0001).

**Supplemental Figure 2:**
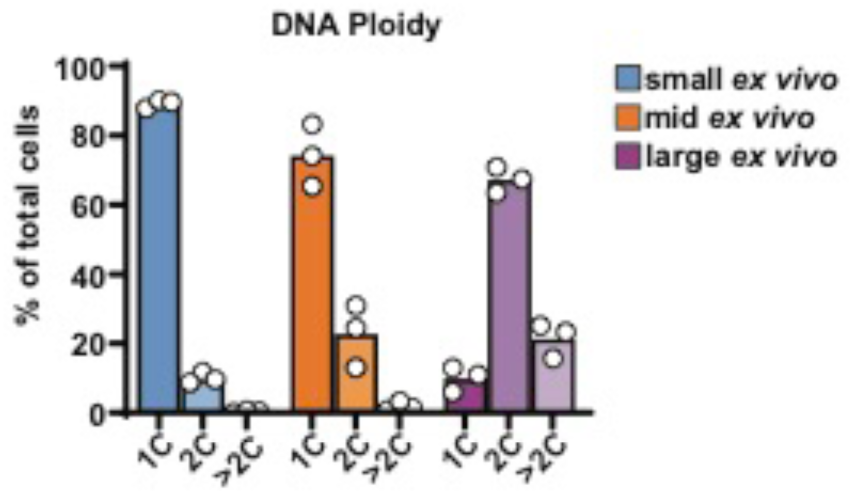
DNA content of *ex vivo C. neoformans* populations. Percentage of cells with 1C, 2C, and >2C DNA content in the indicated population. Values correspond to the representative flow cytometry plot in Figure 2D. Bar graphs display the mean.

**Supplemental Figure 3:**
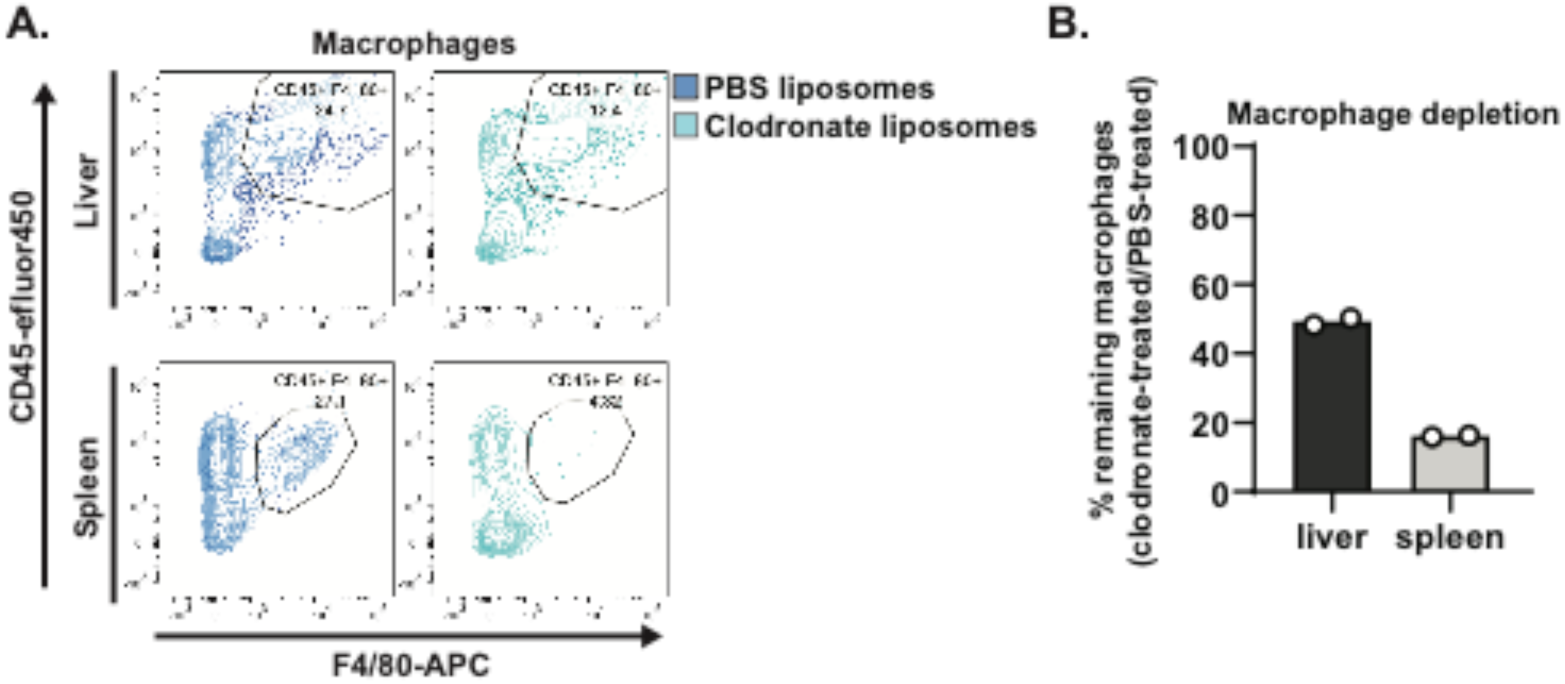
Macrophage depletion efficiency in the liver and spleen. **(A)** Representative flow cytometry plots displaying the frequency of CD45^+^, F4/80^+^ macrophages in the liver and spleen after treatment with either PBS (control) liposomes or clodronate liposomes. **(B)** Quantification of macrophage depletion measured as the ratio of macrophages in clodronate-treated mice to PBS-treated mice.

**Supplemental Figure 4:**
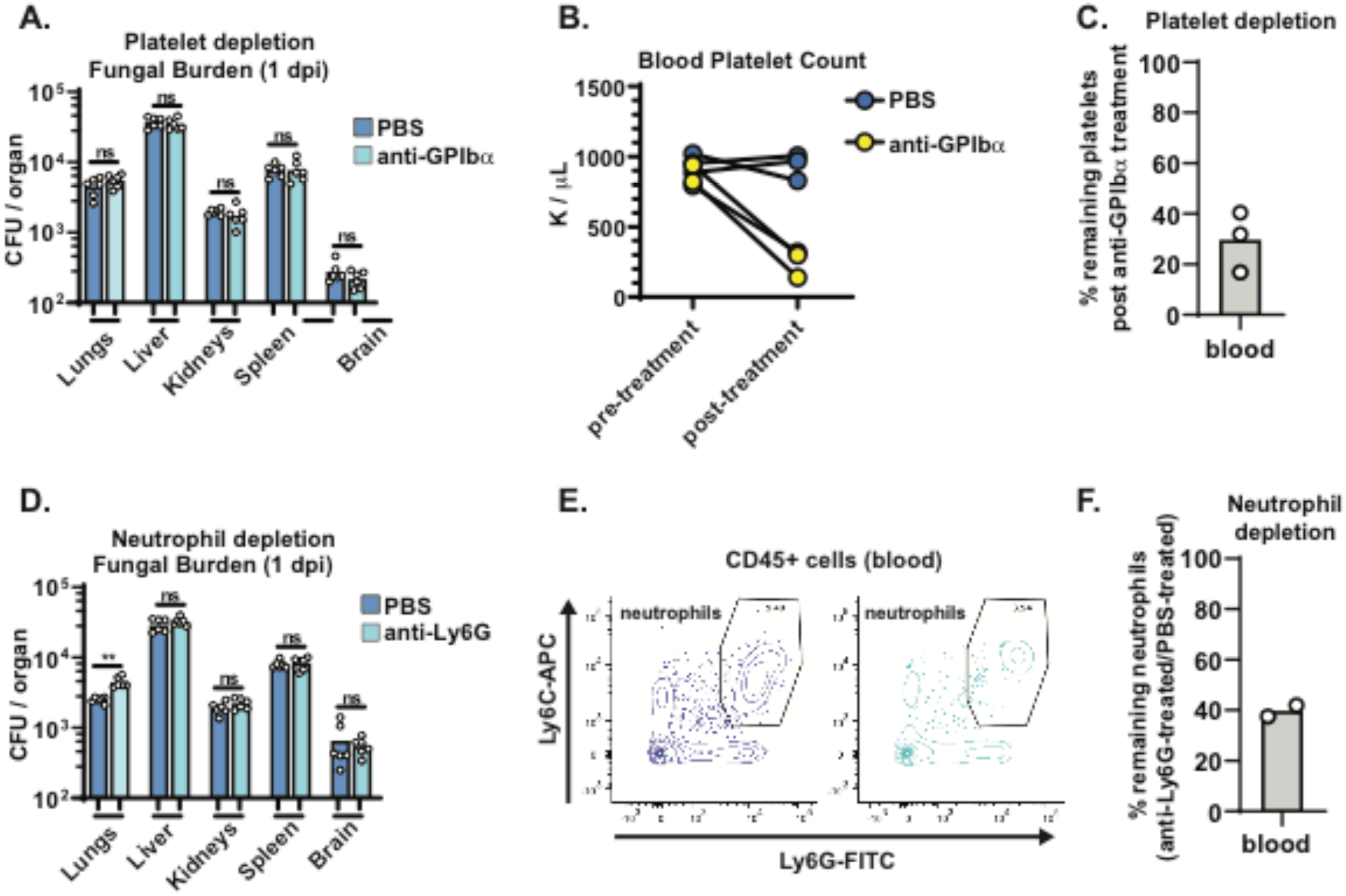
Depletion of platelets or neutrophils does not affect the dissemination of small ex vivo cells. **(A)** Fungal burden in mice, measured by colony forming units (CFU), following treatment with PBS (control) or anti-GPIbα antibody to deplete platelets prior to intravenous inoculation with 10^5^ small *ex vivo* cells / mouse. **(B)** Platelet count (K: units) in the blood before and after treatment with PBS (control) or anti-GPIbα antibody. **(C)** Quantification of platelet depletion in Figure S4B. **(D)** Fungal burden in mice treated with PBS (control) or anti-Ly6G antibody to deplete neutrophils prior to intravenous inoculation with 105 small ex vivo cells / mouse. **(E)** Representative flow cytometry plots displaying the frequency of CD45^+^, Ly6C^+^, Ly6G^+^ neutrophils in the blood after treatment with either PBS (control) or anti- Ly6G antibody. **(F)** Quantification of neutrophil depletion measured as the ratio of neutrophils in anti-Ly6G antibody-treated mice to PBS-treated mice. All comparisons in Figure S4 were analyzed using unpaired t-tests (ns: not significant *p<0.05; **p<0.01; ***p<0.001; ****p<0.0001). All bar graphs display the mean.

**Supplemental Figure 5:**
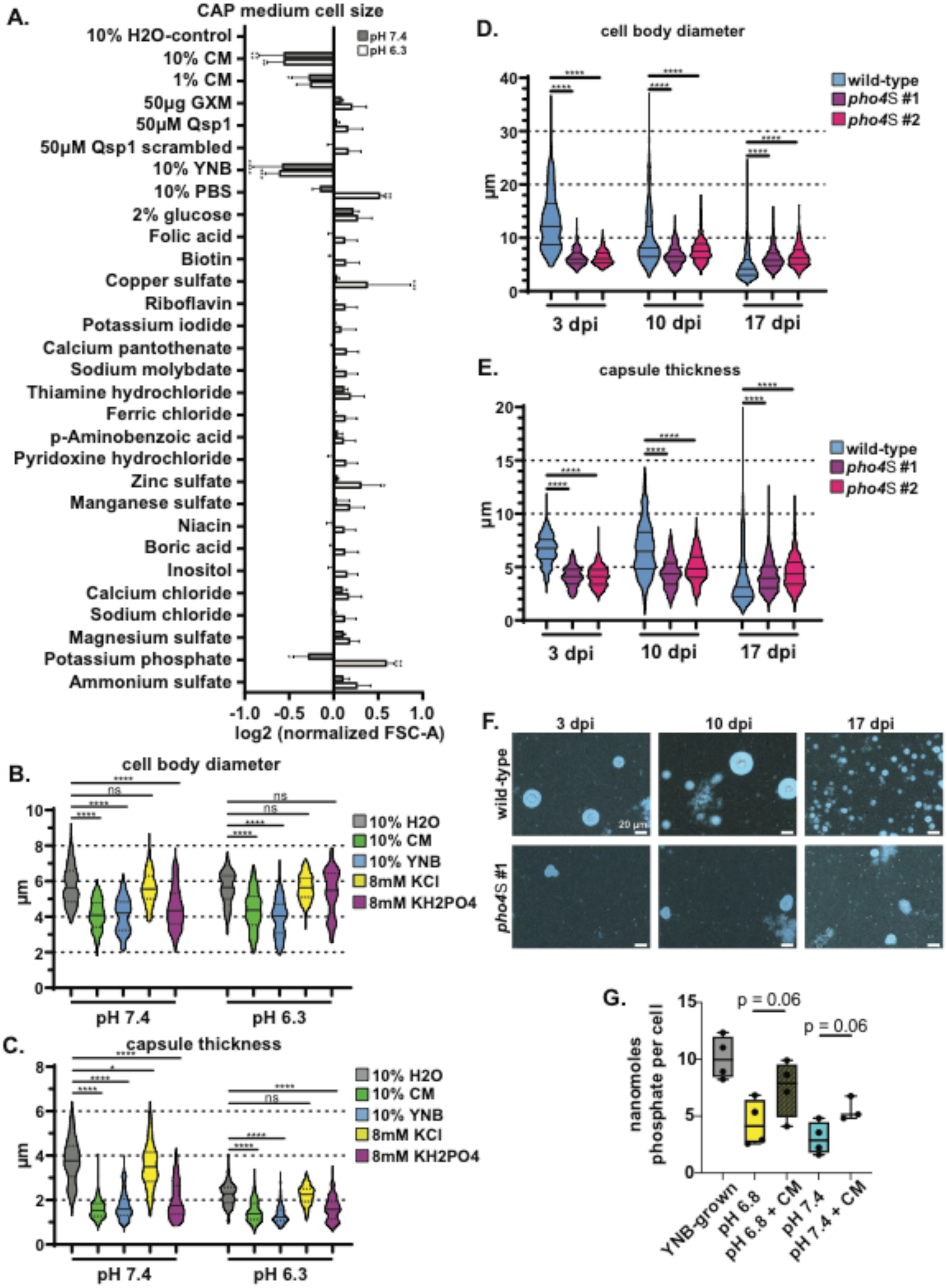
Screen for factors that induce cell size reduction. **(A)** Fungal cell size measured by forward scatter-area (FSC-A). Cells were cultured in capsule inducing medium (CAP medium) buffered to pH 7.4 or 6.3 before subculture (1:1) in fresh medium with 10% of the final volume being the indicated supplement solubilized in H2O (CM: conditioned medium; GXM: glucuronoxylomannan; YNB: minimal YNB medium; PBS: phosphate buffered saline). Comparisons in Figure S5A were analyzed using one-way ANOVA and uncorrected Fisher’s LSD (ns: not significant; *p<0.05; **p<0.01; ***p<0.001; ****p<0.0001). All bar graphs display the mean and error bars indicate standard deviation. **(B)** Cell body diameter and **(C)** capsule thickness of cells cultured in capsule inducing medium (CAP medium) buffered to pH 7.4 or 6.3 before subculture (1:1) in fresh medium with 10% of the final volume being the indicated supplement solubilized in H2O (CM: conditioned medium; YNB: minimal YNB medium). **(D)** Cell body diameter and **(E)** capsule thickness measurements at 3, 10, and 17 days post-intranasal inoculation (dpi) of mice (Mann-Whitney U test; N=6 mice per timepoint; 50, 100, and 150 wild-type cells measured per mouse at days 3, 10, and 17 respectively. 50, 50, and 100 pho4Δ mutant cells measured per mouse at days 3, 10, and 17 respectively). Solid lines in the violin plots indicate the median and dotted lines mark quartiles. All comparisons in Figure S5B-E were analyzed using Mann-Whitney U-tests; N=200 fungal cells (ns: not significant; *p<0.05; **p<0.01; ***p<0.001; ****p<0.0001). **(F)** Representative India ink images of fungal cells quantified in **Fig. 6G**. Scale bars represent 20 μm. (G) Nanomoles of phosphate cells grown in YNB, CAP-medium (pH 6.8 or pH 7.4)

**Supplemental Figure 6:**
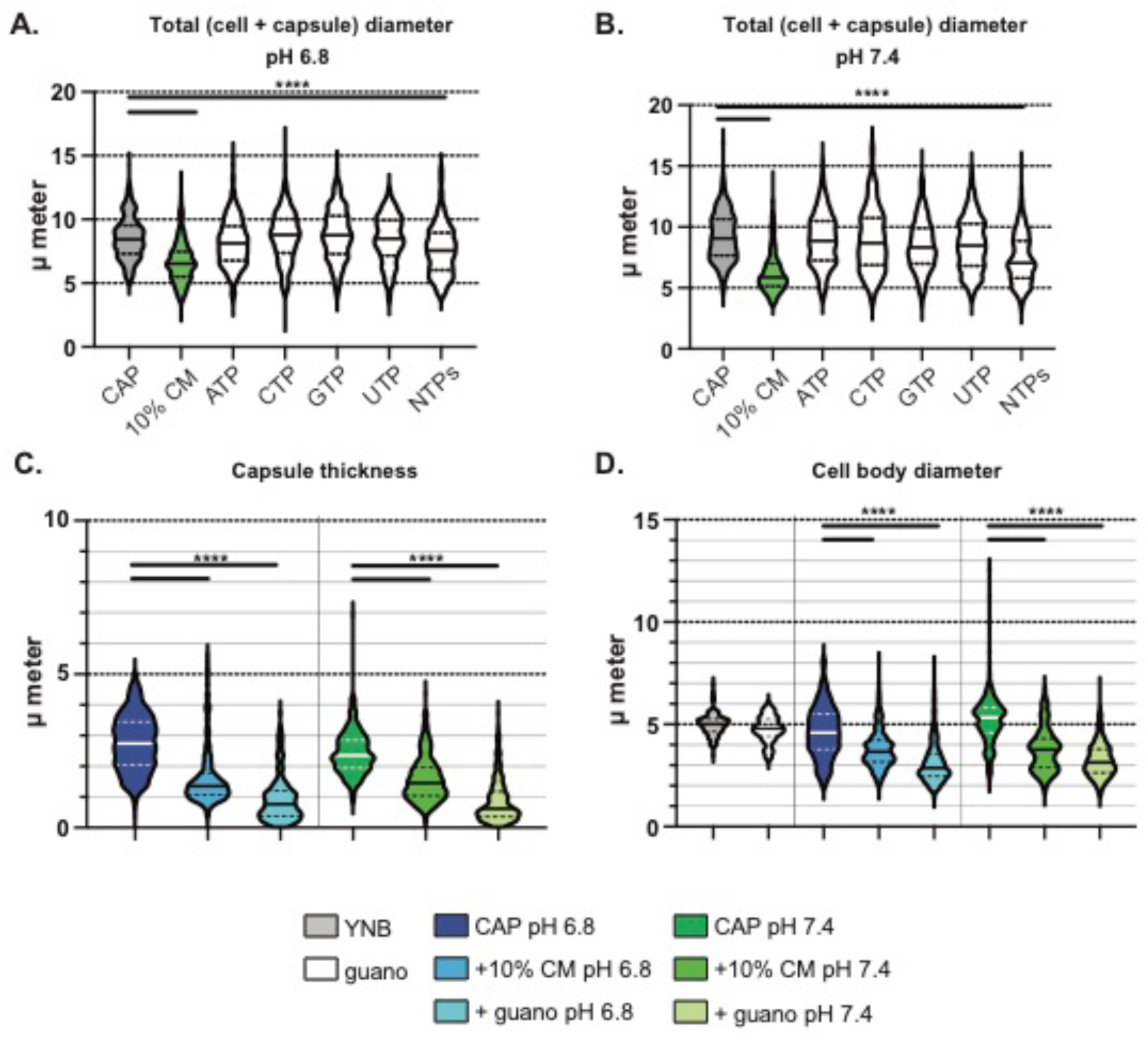
Phosphate is sufficient to induce cell body diameter and capsule thickness reduction in alkaline capsule-inducing medium and is required for morphogenesis *in vivo*. Total diameter measurements of CAP-grown cells (pH 6.8 in A, pH 7.4 in B) after exposure to ATP, CTP, GTP, UTP at 200 μM or a pool of NTPs sy 200 μM total concentration (50 μM each of adenine, cysteine, guanine, and uridine triphosphates). (C) Capsule thickness (calculated using the formula (total diameter – cell body diameter)/2)) of CAP-grown cells (pH 6.8 on left, pH 7.4 on right) induced to form small cells in guano medium. (D) Cell body diameter of CAP-grown cells (pH 6.8 on left, pH 7.4 on right) induced to form small cells in guano medium. P-values calculated using Mann-Whitney test; N = 3 biological replicates per experiment and >100 cells measured per replicate. The solid line represents the population median and the dotted lines the 25^th^ and 75^th^ percentiles. The solid line represents the population median and the dotted lines the 25^th^ and 75^th^ percentiles. All comparisons marked as significant represent populations for which the 95% confidence interval does not overlap. ****p<0.0001.

## REFERENCES

Agustinho, D.P., Miller, L.C., Li, L.X., and Doering, T.L. (2018). Peeling the onion: the outer layers of Cryptococcus neoformans. Mem Inst Oswaldo Cruz 113, e180040.

Aimanianda, V., Bayry, J., Bozza, S., Kniemeyer, O., Perruccio, K., Elluru, S.R., Clavaud, C., Paris, S., Brakhage, A.A., Kaveri, S.V., et al. (2009). Surface hydrophobin prevents immune recognition of airborne fungal spores. Nature 460, 1117–1121.

Alanio, A., Vernel-Pauillac, F., Sturny-Leclère, A., and Dromer, F. (2015a). Cryptococcus neoformans host adaptation: toward biological evidence of dormancy. mBio 6.

Alanio, A., Vernel-Pauillac, F., Sturny-Leclère, A., and Dromer, F. (2015b). *Cryptococcus neoformans* Host Adaptation: Toward Biological Evidence of Dormancy. mBio 6, e02580–02514.

Albuquerque, P., Nicola, A.M., Nieves, E., Paes, H.C., Williamson, P.R., Silva-Pereira, I., and Casadevall, A. (2013). Quorum sensing-mediated, cell density-dependent regulation of growth and virulence in *Cryptococcus neoformans*. mBio 5, e00986–00913.

Altamirano, S., Jackson, K.M., and Nielsen, K. (2020). The interplay of phenotype and genotype in Cryptococcus neoformans disease. Biosci Rep 40, BSR20190337.

André, A.C., Laborde, M., and Marteyn, B.S. (2022). The battle for oxygen during bacterial and fungal infections. Trends Microbiol.

Ballou, E.R., Avelar, G.M., Childers, D.S., Mackie, J., Bain, J.M., Wagener, J., Kastora, S.L., Panea, M.D., Hardison, S.E., Walker, L.A., et al. (2016). Lactate signalling regulates fungal β-glucan masking and immune evasion. Nat Microbiol 2, 16238.

Basenko, E.Y., Pulman, J.A., Shanmugasundram, A., Harb, O.S., Crouch, K., Starns, D., Warrenfeltz, S., Aurrecoechea, C., Stoeckert, C.J., Jr., Kissinger, J.C., et al. (2018). FungiDB: An Integrated Bioinformatic Resource for Fungi and Oomycetes. J Fungi (Basel) 4, 39.

Becquer, A., Trap, J., Irshad, U., Ali, M.A., and Claude, P. (2014). From soil to plant, the journey of P through trophic relationships and ectomycorrhizal association. Front Plant Sci 5, 548.

Beis, I., and Newsholme, E.A. (1975). The contents of adenine nucleotides, phosphagens and some glycolytic intermediates in resting muscles from vertebrates and invertebrates. Biochem J 152, 23–32.

Bojarczuk, A., Miller, K.A., Hotham, R., Lewis, A., Ogryzko, N.V., Kamuyango, A.A., Frost, H., Gibson, R.H., Stillman, E., May, R.C.*, et al.* (2016). *Cryptococcus neoformans* intracellular proliferation and capsule size determines early macrophage control of infection. Sci Rep 6, 21489.

Botts, M.R., Giles, S.S., Gates, M.A., Kozel, T.R., and Hull, C.M. (2009). Isolation and Characterization of *Cryptococcus neoformans* Spores Reveal a Critical Role for Capsule Biosynthesis Genes in Spore Biogenesis. Eukaryot Cell 8, 595–605.

Botts, M.R., and Hull, C.M. (2010). Dueling in the lung: how Cryptococcus spores race the host for survival. Curr Opin Microbiol 13, 437–442.

Brinkmann, V., Reichard, U., Goosmann, C., Fauler, B., Uhlemann, Y., Weiss, D.S., Weinrauch, Y., and Zychlinsky, A. (2004). Neutrophil extracellular traps kill bacteria. Science 303, 1532–1535.

Broadley, S.P., Plaumann, A., Coletti, R., Lehmann, C., Wanisch, A., Seidlmeier, A., Esser, K., Luo, S., Ramer, P.C., Massberg, S., et al. (2016). Dual-Track Clearance of Circulating Bacteria Balances Rapid Restoration of Blood Sterility with Induction of Adaptive Immunity. Cell Host Microbe 20, 36–48.

Brown, G.D., Herre, J., Williams, D.L., Willment, J.A., Marshall, A.S., and Gordon, S. (2003). Dectin-1 mediates the biological effects of beta-glucans. J Exp Med 197, 1119–1124.

Bulmer, G.S., and Sans, M.D. (1967). Cryptococcus neoformans. II. Phagocytosis by human leukocytes. J Bacteriol 94, 1480–1483.

Casadevall, A., Coelho, C., Cordero, R.J.B., Dragotakes, Q., Jung, E., Vij, R., and Wear, M.P. (2019). The capsule of *Cryptococcus neoformans*. Virulence 10, 822–831.

Chaconas, G., Castellanos, M., and Verhey, T.B. (2020). Changing of the guard: How the Lyme disease spirochete subverts the host immune response. The Journal of biological chemistry 295, 301–313.

Chang, Y.C., Stins, M.F., McCaffery, M.J., Miller, G.F., Pare, D.R., Dam, T., Paul-Satyaseela, M., Kim, K.S., and Kwon-Chung, K.J. (2004). Cryptococcal yeast cells invade the central nervous system via transcellular penetration of the blood-brain barrier. Infect Immun 72, 4985–4995.

Charlier, C., Chrétien, F., Baudrimont, M., Mordelet, E., Lortholary, O., and Dromer, F. (2005). Capsule structure changes associated with *Cryptococcus neoformans* crossing of the blood-brain barrier. Am J Pathol 166, 421–432.

Charlier, C., Nielsen, K., Daou, S., Brigitte, M., Chretien, F., and Dromer, F. (2009). Evidence of a role for monocytes in dissemination and brain invasion by Cryptococcus neoformans. Infect Immun 77, 120–127.

Chen, S.H.M., Stins, M.F., Huang, S.H., Chen, Y.H., Kwon-Chung, K.J., Chang, Y., Kim, K.S., Suzuki, K., and Jong, A.Y. (2003). Cryptococcus neoformans induces alterations in the cytoskeleton of human brain microvascular endothelial cells. J Med Microbiol 52, 961–970.

Chen, Y., Li, C., Sun, D., Strickland, A.B., Liu, G., and Shi, M. (2021). Quantitative analysis reveals internalisation of Cryptococcus neoformans by brain endothelial cells in vivo. Cell Microbiol 23, e13330.

Chun, C.D., Brown, J.C.S., and Madhani, H.D. (2011). A Major Role for Capsule-Independent Phagocytosis-Inhibitory Mechanisms in Mammalian Infection by *Cryptococcus neoformans*. Cell Host Microbe 9, 243–251.

Chung, K.Y., and Brown, J.C.S. (2020). Biology and function of exo-polysaccharides from human fungal pathogens. Curr Clin Microbiol Rep 7, 1–11.

Crabtree, J.N., Okagaki, L.H., Wiesner, D.L., Strain, A.K., Nielsen, J.N., and Nielsen, K. (2012). Titan cell production enhances the virulence of Cryptococcus neoformans. Infect Immun 80, 3776–3785.

Dambuza, I.M., Drake, T., Chapuis, A., Zhou, X., Correia, J., Taylor-Smith, L., LeGrave, N., Rasmussen, T., Fisher, M.C., Bicanic, T., et al. (2018). The Cryptococcus neoformans Titan cell is an inducible and regulated morphotype underlying pathogenesis. PLoS pathogens 14, e1006978–e1006978.

Dan, J.M., Kelly, R.M., Lee, C.K., and Levitz, S.M. (2008). Role of the mannose receptor in a murine model of *Cryptococcus neoformans* infection. Infect Immun 76, 2362–2367.

Davidson, R.C., Cruz, M.C., Sia, R.A., Allen, B., Alspaugh, J.A., and Heitman, J. (2000). Gene disruption by biolistic transformation in serotype D strains of Cryptococcus neoformans. Fungal Genet Biol 29, 38–48.

Davis, K.M. (2018). All Yersinia Are Not Created Equal: Phenotypic Adaptation to Distinct Niches Within Mammalian Tissues. Front Cell Infect Microbiol 8, 261.

Davis, K.M., Mohammadi, S., and Isberg, R.R. (2015). Community behavior and spatial regulation within a bacterial microcolony in deep tissue sites serves to protect against host attack. Cell Host Microbe 17, 21–31.

Deitsch, K.W., del Pinal, A., and Wellems, T.E. (1999). Intra-cluster recombination and var transcription switches in the antigenic variation of Plasmodium falciparum. Mol Biochem Parasitol 101, 107–116.

Denham, S.T., and Brown, J.C.S. (2018). Mechanisms of Pulmonary Escape and Dissemination by Cryptococcus neoformans. J Fungi (Basel) 4, 25.

Denham, S.T., Verma, S., Reynolds, R.C., Worne, C.L., Daugherty, J.M., Lane, T.E., and Brown, J.C.S. (2018). Regulated release of cryptococcal polysaccharide drives virulence and suppresses immune infiltration into the central nervous system. Infect Immun 86.

Diamond, R.D., and Bennett, J.E. (1973). Growth of Cryptococcus neoformans Within Human Macrophages In Vitro. Infect Immun 7, 231–236.

Dobin, A., Davis, C.A., Schlesinger, F., Drenkow, J., Zaleski, C., Jha, S., Batut, P., Chaisson, M., and Gingeras, T.R. (2013). STAR: ultrafast universal RNA-seq aligner. Bioinformatics 29, 15–21.

Dosch, M., Gerber, J., Jebbawi, F., and Beldi, G. (2018). Mechanisms of ATP Release by Inflammatory Cells. Int J Mol Sci 19.

Eisen, D.P., Dean, M.M., O’Sullivan, M.V., Heatley, S., and Minchinton, R.M. (2008). Mannose-binding lectin deficiency does not appear to predispose to cryptococcosis in non-immunocompromised patients. Med Mycol 46, 371–375.

Elliott, M.R., Chekeni, F.B., Trampont, P.C., Lazarowski, E.R., Kadl, A., Walk, S.F., Park, D., Woodson, R.I., Ostankovich, M., Sharma, P., et al. (2009). Nucleotides released by apoptotic cells act as a find-me signal to promote phagocytic clearance. Nature 461, 282–286.

Emmons, C.W. (1955). Saprophytic sources of Cryptococcus neoformans associated with the pigeon (Columba livia). Am J Hyg 62, 227–232.

Ewels, P., Magnusson, M., Lundin, S., and Kaller, M. (2016). MultiQC: summarize analysis results for multiple tools and samples in a single report. Bioinformatics 32, 3047–3048.

Fang, W., Fa, Z., and Liao, W. (2015). Epidemiology of *Cryptococcus* and cryptococcosis in China. Fungal Genet Biol 78, 7–15.

Farhi, F., Bulmer, G.S., and Tacker, J.R. (1970). Cryptococcus neoformans IV. The Not-So-Encapsulated Yeast. Infect Immun 1, 526–531.

Feldmesser, M., Kress, Y., and Casadevall, A. (2001). Dynamic changes in the morphology of Cryptococcus neoformans during murine pulmonary infection. Microbiology 147, 2355–2365.

Feldmesser, M., Kress, Y., Novikoff, P., and Casadevall, A. (2000). *Cryptococcus neoformans* Is a facultative intracellular pathogen in murine pulmonary infection. Infect Immun 68, 4225–4237.

Fernandes, K.E., Brockway, A., Haverkamp, M., Cuomo, C.A., van Ogtrop, F., Perfect, J.R., and Carter, D.A. (2018). Phenotypic Variability Correlates with Clinical Outcome in Cryptococcus Isolates Obtained from Botswanan HIV/AIDS Patients. mBio 9.

Fernandes, K.E., and Carter, D.A. (2020). Cellular plasticity of pathogenic fungi during infection. PLoS Pathog 16, e1008571.

Fernandes, K.E., Dwyer, C., Campbell, L.T., and Carter, D.A. (2016). Species in the Cryptococcus gattii Complex Differ in Capsule and Cell Size following Growth under Capsule-Inducing Conditions. mSphere 1.

Garfoot, A.L., Shen, Q., Wüthrich, M., Klein, B.S., and Rappleye, C.A. (2016). The Eng1 β-Glucanase Enhances Histoplasma Virulence by Reducing β-Glucan Exposure. mBio 7, e01388–01315.

Gault, W.J., Enyedi, B., and Niethammer, P. (2014). Osmotic surveillance mediates rapid wound closure through nucleotide release. J Cell Biol 207, 767–782.

Gaylord, E.A., Choy, H.L., and Doering, T.L. (2020). Dangerous Liaisons: Interactions of Cryptococcus neoformans with Host Phagocytes. Pathogens 9.

Gerstein, A.C., Fu, M.S., Mukaremera, L., Li, Z., Ormerod, K.L., Fraser, J.A., Berman, J., and Nielsen, K. (2015). Polyploid titan cells produce haploid and aneuploid progeny to promote stress adaptation. mBio 6, e01340.

Geunes-Boyer, S., Beers, M.F., Perfect, J.R., Heitman, J., and Wright, J.R. (2012). Surfactant protein D facilitates *Cryptococcus neoformans* infection. Infect Immun 80, 2444–2453.

Goldman, D.L., Khine, H., Abadi, J., Lindenberg, D.J., Pirofski, L., Niang, R., and Casadevall, (2001). Serologic evidence for *Cryptococcus neoformans* infection in early childhood. Pediatrics 107, E66.

Gow, N.A., Netea, M.G., Munro, C.A., Ferwerda, G., Bates, S., Mora-Montes, H.M., Walker, L., Jansen, T., Jacobs, L., Tsoni, V.*, et al.* (2007). Immune recognition of Candida albicans beta-glucan by dectin-1. J Infect Dis 196, 1565–1571.

Gow, N.A.R., Latge, J.P., and Munro, C.A. (2017). The Fungal Cell Wall: Structure, Biosynthesis, and Function. Microbiol Spectr 5.

Grygorczyk, R., Boudreault, F., Ponomarchuk, O., Tan, J.J., Furuya, K., Goldgewicht, J., Kenfack, F.D., and Yu, F. (2021). Lytic Release of Cellular ATP: Physiological Relevance and Therapeutic Applications. Life (Basel) 11.

Guimarães, A.J., de Cerqueira, M.D., and Nosanchuk, J.D. (2011). Surface architecture of histoplasma capsulatum. Front Microbiol 2, 225.

Haas, R., and Meyer, T.F. (1986). The repertoire of silent pilus genes in Neisseria gonorrhoeae: evidence for gene conversion. Cell 44, 107–115.

Harris, G., Davies, J.W., and Parsons, R. (1958). Nucleotide pool of brewers’ yeast during growth and fermentation. Nature 182, 1565–1567.

Hasan, M.M., Hasan, M.M., Teixeira da Silva, J.A., and Li, X. (2016). Regulation of phosphorus uptake and utilization: transitioning from current knowledge to practical strategies. Cell Mol Biol Lett 21, 7.

Himmelreich, U., Allen, C., Dowd, S., Malik, R., Shehan, B.P., Mountford, C., and Sorrell, T.C. (2003). Identification of metabolites of importance in the pathogenesis of pulmonary cryptococcoma using nuclear magnetic resonance spectroscopy. Microbes and Infection 5, 285–290.

Hohl, T.M., Van Epps, H.L., Rivera, A., Morgan, L.A., Chen, P.L., Feldmesser, M., and Pamer, E.G. (2005). Aspergillus fumigatus triggers inflammatory responses by stage-specific beta-glucan display. PLoS Pathog 1, e30.

Homer, C.M., Summers, D.K., Goranov, A.I., Clarke, S.C., Wiesner, D.L., Diedrich, J.K., Moresco, J.J., Toffaletti, D., Upadhya, R., Caradonna, I.*, et al.* (2016). Intracellular Action of a Secreted Peptide Required for Fungal Virulence. Cell Host Microbe 19, 849–864.

Hommel, B., Mukaremera, L., Cordero, R.J.B., Coelho, C., Desjardins, C.A., Sturny-Leclère, A., Janbon, G., Perfect, J.R., Fraser, J.A., Casadevall, A.*, et al.* (2018). Titan cells formation in Cryptococcus neoformans is finely tuned by environmental conditions and modulated by positive and negative genetic regulators. PLoS Pathog 14, e1006982.

Hood, M.I., and Skaar, E.P. (2012). Nutritional immunity: transition metals at the pathogen-host interface. Nature Reviews Microbiology 10, 525–537.

Huang, S.H., Long, M., Wu, C.H., Kwon-Chung, K.J., Chang, Y.C., Chi, F., Lee, S., and Jong, A. (2011). Invasion of Cryptococcus neoformans into human brain microvascular endothelial cells is mediated through the lipid rafts-endocytic pathway via the dual specificity tyrosine phosphorylation-regulated kinase 3 (DYRK3). J Biol Chem 286, 34761–34769.

Ikeh, M.A., Kastora, S.L., Day, A.M., Herrero-de-Dios, C.M., Tarrant, E., Waldron, K.J., Banks, A.P., Bain, J.M., Lydall, D., Veal, E.A.*, et al.* (2016). Pho4 mediates phosphate acquisition in *Candida albicans* and is vital for stress resistance and metal homeostasis. Mol Biol Cell 27, 2784–2801.

Jacobs, C.W., Adams, A.E., Szaniszlo, P.J., and Pringle, J.R. (1988). Functions of microtubules in the Saccharomyces cerevisiae cell cycle. J Cell Biol 107, 1409–1426.

Kanetsuna, F., and Carbonell, L.M. (1971). Cell wall composition of the yeastlike and mycelial forms of Blastomyces dermatitidis. J Bacteriol 106, 946–948.

Kim, K. (2012). Malaria var gene expression: keeping up with the neighbors. Cell Host Microbe 11, 1–2.

Kohler, J.R., Acosta-Zaldivar, M., and Qi, W. (2020). Phosphate in Virulence of Candida albicans and Candida glabrata. J Fungi (Basel) 6, 40.

Kowalski, C.H., Kerkaert, J.D., Liu, K.W., Bond, M.C., Hartmann, R., Nadell, C.D., Stajich, J.E., and Cramer, R.A. (2019). Fungal biofilm morphology impacts hypoxia fitness and disease progression. Nat Microbiol 4, 2430–2441.

Kretschmer, M., Reiner, E., Hu, G., Tam, N., Oliveira, D.L., Caza, M., Yeon, J.H., Kim, J., Kastrup, C.J., Jung, W.H., et al. (2014). Defects in phosphate acquisition and storage influence virulence of Cryptococcus neoformans. Infect Immun 82, 2697–2712.

Kubes, P., and Jenne, C. (2018). Immune Responses in the Liver. Annu Rev Immunol 36, 247–277.

Lazera, M.S., Pires, F.D., Camillo-Coura, L., Nishikawa, M.M., Bezerra, C.C., Trilles, L., and Wanke, B. (1996). Natural habitat of *Cryptococcus neoformans var. neoformans* in decaying wood forming hollows in living trees. J Med Vet Mycol 34, 127–131.

Lee, H., Chang, Y.C., Nardone, G., and Kwon-Chung, K.J. (2007). TUP1 disruption in *Cryptococcus neoformans* uncovers a peptide-mediated density-dependent growth phenomenon that mimics quorum sensing. Mol Microbiol 64, 591–601.

Lee, S.C., Dickson, D.W., and Casadevall, A. (1996). Pathology of cryptococcal meningoencephalitis: analysis of 27 patients with pathogenetic implications. Hum Pathol 27, 839–847.

Lev, S., and Djordjevic, J.T. (2018). Why is a functional PHO pathway required by fungal pathogens to disseminate within a phosphate-rich host: A paradox explained by alkaline pH-simulated nutrient deprivation and expanded PHO pathway function. PLoS Pathog 14, e1007021.

Lev, S., Kaufman-Francis, K., Desmarini, D., Juillard, P.G., Li, C., Stifter, S.A., Feng, C.G., Sorrell, T.C., Grau, G.E., Bahn, Y.S., et al. (2017). Pho4 Is Essential for Dissemination of Cryptococcus neoformans to the Host Brain by Promoting Phosphate Uptake and Growth at Alkaline pH. mSphere 2.

Li, P.Z., Li, J.Z., Li, M., Gong, J.P., and He, K. (2014). An efficient method to isolate and culture mouse Kupffer cells. Immunol Lett 158, 52–56.

Li, Z., and Nielsen, K. (2017). Morphology Changes in Human Fungal Pathogens upon Interaction with the Host. J Fungi (Basel) 3, 66.

Liao, Y., Smyth, G.K., and Shi, W. (2014). featureCounts: an efficient general purpose program for assigning sequence reads to genomic features. Bioinformatics 30, 923–930.

Lim, J., Coates, C.J., Seoane, P.I., Garelnabi, M., Taylor-Smith, L.M., Monteith, P., Macleod, C.L., Escaron, C.J., Brown, G.D., Hall, R.A.*, et al.* (2018). Characterizing the Mechanisms of Nonopsonic Uptake of Cryptococci by Macrophages. J Immunol 200, 3539–3546.

Lin, Y.Y., Shiau, S., and Fang, C.T. (2015). Risk factors for invasive *Cryptococcus neoformans* diseases: a case-control study. PLoS One 10, e0119090.

Liu, N.N., Uppuluri, P., Broggi, A., Besold, A., Ryman, K., Kambara, H., Solis, N., Lorenz, V., Qi, W., Acosta-Zaldivar, M., et al. (2018). Intersection of phosphate transport, oxidative stress and TOR signalling in *Candida albicans* virulence. PLoS Pathog 14, e1007076.

Liu, T.B., Kim, J.C., Wang, Y., Toffaletti, D.L., Eugenin, E., Perfect, J.R., Kim, K.J., and Xue, C. (2013). Brain inositol is a novel stimulator for promoting Cryptococcus penetration of the blood-brain barrier. PLoS Pathog 9, e1003247.

Liu, T.B., Subbian, S., Pan, W., Eugenin, E., Xie, J., and Xue, C. (2014). Cryptococcus inositol utilization modulates the host protective immune response during brain infection. Cell Communication and Signaling 12, 51.

López-Fuentes, E., Gutiérrez-Escobedo, G., Timmermans, B., Van Dijck, P., De Las Peñas, A., and Castaño, I. (2018). Candida glabrata’s Genome Plasticity Confers a Unique Pattern of Expressed Cell Wall Proteins. J Fungi (Basel) 4.

Love, M.I., Huber, W., and Anders, S. (2014). Moderated estimation of fold change and dispersion for RNA-seq data with DESeq2. Genome Biol 15, 550.

Luberto, C., Martinez-Marino, B., Taraskiewicz, D., Bolanos, B., Chitano, P., Toffaletti, D.L., Cox, G.M., Perfect, J.R., Hannun, Y.A., Balish, E., et al. (2003). Identification of App1 as a regulator of phagocytosis and virulence of *Cryptococcus neoformans*. J Clin Invest 112, 1080–1094.

Macpherson, A.J., Heikenwalder, M., and Ganal-Vonarburg, S.C. (2016). The Liver at the Nexus of Host-Microbial Interactions. Cell Host Microbe 20, 561–571.

Martin, M. (2011). Cutadapt removes adapter sequences from high-throughput sequencing reads. EMBnet 17, 10–12.

Maruvada, R., Zhu, L., Pearce, D., Zheng, Y., Perfect, J., Kwon-Chung, K.J., and Kim, K.S. (2012). Cryptococcus neoformans phospholipase B1 activates host cell Rac1 for traversal across the blood-brain barrier. Cell Microbiol 14, 1544–1553.

Maziarz, E.K., and Perfect, J.R. (2016). Cryptococcosis. Infect Dis Clin North Am 30, 179–206.

Mukaremera, L., Lee, K.K., Mora-Montes, H.M., and Gow, N.A.R. (2017). Candida albicans Yeast, Pseudohyphal, and Hyphal Morphogenesis Differentially Affects Immune Recognition. Front Immunol 8, 629.

Mukaremera, L., McDonald, T.R., Nielsen, J.N., Molenaar, C.J., Akampurira, A., Schutz, C., Taseera, K., Muzoora, C., Meintjes, G., Meya, D.B.*, et al.* (2019). The Mouse Inhalation Model of Cryptococcus neoformans Infection Recapitulates Strain Virulence in Humans and Shows that Closely Related Strains Can Possess Differential Virulence. Infect Immun 87, e00046–00019.

Ngamskulrungroj, P., Chang, Y., Sionov, E., and Kwon-Chung, K.J. (2012). The primary target organ of *Cryptococcus gattii* is different from that of *Cryptococcus neoformans* in a murine model. mBio 3, e00103–00112.

O’Meara, T.R., and Alspaugh, J.A. (2012). The *Cryptococcus neoformans* capsule: a sword and a shield. Clin Microbiol Rev 25, 387–408.

Okagaki, L.H., and Nielsen, K. (2012). Titan cells confer protection from phagocytosis in Cryptococcus neoformans infections. Eukaryot Cell 11, 820–826.

Okagaki, L.H., Strain, A.K., Nielsen, J.N., Charlier, C., Baltes, N.J., Chrétien, F., Heitman, J., Dromer, F., and Nielsen, K. (2010). Cryptococcal cell morphology affects host cell interactions and pathogenicity. PLoS Pathog 6, e1000953–e1000953.

Okagaki, L.H., Wang, Y., Ballou, E.R., O’Meara, T.R., Bahn, Y.-S., Alspaugh, J.A., Xue, C., and Nielsen, K. (2011). Cryptococcal titan cell formation is regulated by G-protein signaling in response to multiple stimuli. Eukaryot Cell 10, 1306–1316.

Olszewski, M.A., Noverr, M.C., Chen, G.H., Toews, G.B., Cox, G.M., Perfect, J.R., and Huffnagle, G.B. (2004). Urease expression by Cryptococcus neoformans promotes microvascular sequestration, thereby enhancing central nervous system invasion. Am J Pathol 164, 1761–1771.

Otero, X.L., De La Peña-Lastra, S., Pérez-Alberti, A., Ferreira, T.O., and Huerta-Diaz, M.A. (2018). Seabird colonies as important global drivers in the nitrogen and phosphorus cycles. Nat Commun 9, 246.

Pande, K., Chen, C., and Noble, S.M. (2013). Passage through the mammalian gut triggers a phenotypic switch that promotes Candida albicans commensalism. Nat Genet 45, 1088–1091.

Panepinto, J.C., Komperda, K.W., Hacham, M., Shin, S., Liu, X., and Williamson, P.R. (2007). Binding of serum mannan binding lectin to a cell integrity-defective *Cryptococcus neoformans* ccr4Delta mutant. Infect Immun 75, 4769–4779.

Pearson, A.M., Rich, A., and Krieger, M. (1993). Polynucleotide binding to macrophage scavenger receptors depends on the formation of base-quartet-stabilized four-stranded helices. J Biol Chem 268, 3546–3554.

Rajasingham, R., Smith, R.M., Park, B.J., Jarvis, J.N., Govender, N.P., Chiller, T.M., Denning, D.W., Loyse, A., and Boulware, D.R. (2017). Global burden of disease of HIV-associated cryptococcal meningitis: an updated analysis. Lancet Infect Dis 17, 873–881.

Ren, H., Teng, Y., Tan, B., Zhang, X., Jiang, W., Liu, M., Jiang, W., Du, B., and Qian, M. (2014). Toll-like receptor-triggered calcium mobilization protects mice against bacterial infection through extracellular ATP release. Infect Immun 82, 5076–5085.

Rivera, J., Feldmesser, M., Cammer, M., and Casadevall, A. (1998). Organ-Dependent Variation of Capsule Thickness in Cryptococcus neoformans during Experimental Murine Infection. Infect Immun 66, 5027–5030.

Roberts, D.J., Craig, A.G., Berendt, A.R., Pinches, R., Nash, G., Marsh, K., and Newbold, C.I. (1992). Rapid switching to multiple antigenic and adhesive phenotypes in malaria. Nature 357, 689–692.

Sabiiti, W., May, R.C., and Pursall, E.R. (2012). Experimental models of cryptococcosis. Int J Microbiol 2012, 626745.

Sahly, H., Ofek, I., Podschun, R., Brade, H., He, Y., Ullmann, U., and Crouch, E. (2002). Surfactant protein D binds selectively to Klebsiella pneumoniae lipopolysaccharides containing mannose-rich O-antigens. J Immunol 169, 3267–3274.

San Juan, B.P., Garcia-Leon, M.J., Rangel, L., Goetz, J.G., and Chaffer, C.L. (2019). The Complexities of Metastasis. Cancers (Basel) 11, 1575.

Santiago-Tirado, F.H., Onken, M.D., Cooper, J.A., Klein, R.S., and Doering, T.L. (2017). Trojan Horse Transit Contributes to Blood-Brain Barrier Crossing of a Eukaryotic Pathogen. mBio 8.

Selmecki, A.M., Maruvka, Y.E., Richmond, P.A., Guillet, M., Shoresh, N., Sorenson, A.L., De, S., Kishony, R., Michor, F., Dowell, R.*, et al.* (2015). Polyploidy can drive rapid adaptation in yeast. Nature 519, 349–352.

Smith, J.D., Chitnis, C.E., Craig, A.G., Roberts, D.J., Hudson-Taylor, D.E., Peterson, D.S., Pinches, R., Newbold, C.I., and Miller, L.H. (1995). Switches in expression of Plasmodium falciparum var genes correlate with changes in antigenic and cytoadherent phenotypes of infected erythrocytes. Cell 82, 101–110.

Stajich, J.E., Harris, T., Brunk, B.P., Brestelli, J., Fischer, S., Harb, O.S., Kissinger, J.C., Li, W., Nayak, V., Pinney, D.F.*, et al.* (2012). FungiDB: an integrated functional genomics database for fungi. Nucleic Acids Res 40, D675–681.

Stano, P., Williams, V., Villani, M., Cymbalyuk, E.S., Qureshi, A., Huang, Y., Morace, G., Luberto, C., Tomlinson, S., and Del Poeta, M. (2009). App1: an antiphagocytic protein that binds to complement receptors 3 and 2. J Immunol 182, 84–91.

Sun, D., Sun, P., Li, H., Zhang, M., Liu, G., Strickland, A.B., Chen, Y., Fu, Y., Xu, J., Yosri, M.*, et al.* (2019). Fungal dissemination is limited by liver macrophage filtration of the blood. Nat Commun 10, 4566.

Sun, D., Zhang, M., Liu, G., Wu, H., Li, C., Zhou, H., Zhang, X., and Shi, M. (2016). Intravascular clearance of disseminating *Cryptococcus neoformans* in the brain can be improved by enhancing neutrophil recruitment in mice. Eur J Immunol 46, 1704–1714.

Swanson, J., Bergström, S., Robbins, K., Barrera, O., Corwin, D., and Koomey, J.M. (1986). Gene conversion involving the pilin structural gene correlates with pilus+ in equilibrium with pilus-changes in Neisseria gonorrhoeae. Cell 47, 267–276.

Tao, L., Du, H., Guan, G., Dai, Y., Nobile, C.J., Liang, W., Cao, C., Zhang, Q., Zhong, J., and Huang, G. (2014). Discovery of a "white-gray-opaque" tristable phenotypic switching system in candida albicans: roles of non-genetic diversity in host adaptation. PLoS Biol 12, e1001830.

Todd, R.T., Braverman, A.L., and Selmecki, A. (2018). Flow Cytometry Analysis of Fungal Ploidy. Curr Protoc Microbiol 50, e58–e58.

Trevijano-Contador, N., de Oliveira, H.C., García-Rodas, R., Rossi, S.A., Llorente, I., Zaballos, Á., Janbon, G., Ariño, J., and Zaragoza, Ó. (2018). Cryptococcus neoformans can form titan-like cells in vitro in response to multiple signals. PLoS Pathog 14, e1007007.

van der Woude, M.W., and Bäumler, A.J. (2004). Phase and antigenic variation in bacteria. Clin Microbiol Rev 17, 581–611, table of contents.

Viriyakosol, S., Singhania, A., Fierer, J., Goldberg, J., Kirkland, T.N., and Woelk, C.H. (2013). Gene expression in human fungal pathogen Coccidioides immitis changes as arthroconidia differentiate into spherules and mature. BMC Microbiol 13, 121.

Vu, K., Eigenheer, R.A., Phinney, B.S., and Gelli, A. (2013). Cryptococcus neoformans promotes its transmigration into the central nervous system by inducing molecular and cellular changes in brain endothelial cells. Infect Immun 81, 3139–3147.

Walsh, N.M., Botts, M.R., McDermott, A.J., Ortiz, S.C., Wüthrich, M., Klein, B., and Hull, C.M. (2019). Infectious particle identity determines dissemination and disease outcome for the inhaled human fungal pathogen Cryptococcus. PLoS Pathog 15, e1007777.

Wang, L., and Lin, X. (2015). The morphotype heterogeneity in *Cryptococcus neoformans*. Curr Opin Microbiol 26, 60–64.

Wang, L., Zhai, B., and Lin, X. (2012). The link between morphotype transition and virulence in *Cryptococcus neoformans*. PLoS Pathog 8, e1002765.

Ward, C.P., Clottey, G.T., Dorris, M., Ji, D.D., and Arnot, D.E. (1999). Analysis of Plasmodium falciparum PfEMP-1/var genes suggests that recombination rearranges constrained sequences. Mol Biochem Parasitol 102, 167–177.

Weigel, W.A., and Dersch, P. (2018). Phenotypic heterogeneity: a bacterial virulence strategy. Microbes Infect 20, 570–577.

Whiston, E., Zhang Wise, H., Sharpton, T.J., Jui, G., Cole, G.T., and Taylor, J.W. (2012). Comparative transcriptomics of the saprobic and parasitic growth phases in Coccidioides spp. PLoS One 7, e41034.

Wozniak, K.L., and Levitz, S.M. (2009). Isolation and purification of antigenic components of *Cryptococcus*. Methods Mol Biol 470, 71–83.

Wright, L., Bubb, W., Davidson, J., Santangelo, R., Krockenberger, M., Himmelreich, U., and Sorrell, T. (2002). Metabolites released by *Cryptococcus neoformans var. neoformans* and *var. gattii* differentially affect human neutrophil function. Microbes Infect 4, 1427–1438.

Xie, S., Sao, R., Braun, A., and Bottone, E.J. (2012). Difference in Cryptococcus neoformans cellular and capsule size in sequential pulmonary and meningeal infection: a postmortem study. Diagnostic Microbiology and Infectious Disease 73, 49–52.

Yoneda, A., and Doering, T.L. (2006). A eukaryotic capsular polysaccharide is synthesized intracellularly and secreted via exocytosis. Mol Biol Cell 17, 5131–5140.

Zaragoza, O. (2011). Multiple Disguises for the Same Party: The Concepts of Morphogenesis and Phenotypic Variations in Cryptococcus neoformans. Front Microbiol 2, 181.

Zaragoza, O., García-Rodas, R., Nosanchuk, J.D., Cuenca-Estrella, M., Rodríguez-Tudela, J.L., and Casadevall, A. (2010). Fungal cell gigantism during mammalian infection. PLoS Pathog 6, e1000945–e1000945.

Zhang, M., Sun, D., Liu, G., Wu, H., Zhou, H., and Shi, M. (2016). Real-time *in vivo* imaging reveals the ability of neutrophils to remove *Cryptococcus neoformans* directly from the brain vasculature. J Leukoc Biol 99, 467–473.

Zhou, X., and Ballou, E.R. (2018). The Cryptococcus neoformans Titan Cell: From In Vivo Phenomenon to In Vitro Model. Current Clinical Microbiology Reports 5, 252–260.

Zhou, X., Zafar, H., Sephton-Clark, P., Mohamed, S.H., Chapuis, A., Makarova, M., MacCallum, D.M., Drummond, R.A., Dambuza, I.M., and Ballou, E.R. (2020). Host environmental conditions induce small fungal cell size and alter population heterogeneity in Cryptococcus neoformans. bioRxiv, 2020.2001.2003.894709.

